# Vertical transmission of sponge microbiota is inconsistent and unfaithful

**DOI:** 10.1101/425009

**Authors:** Johannes R. Björk, Cristina Diéz-Vives, Carmen Astudillo-García, Elizabeth A. Archie, José M. Montoya

**Affiliations:** Department of Biological Sciences, University of Notre Dame, United States; Theoretical and Experimental Ecology Station, CNRS-University Paul Sabatier, Moulis, France; Natural History Museum, London, United Kingdom; School of Biological Sciences, University of Auckland, New Zealand

## Abstract

Classic coevolutionary theory predicts that if beneficial microbial symbionts improve host fitness, they should be faithfully transmitted to offspring. More recently, the hologenome theory of evolution predicts resemblance between parent and offspring microbiomes and high partner fidelity between host species and their vertically transmitted microbes. Here, we test these ideas for the first time in multiple coexisting host species with highly diverse microbiota, leveraging known parent-offspring pairs sampled from eight species of wild marine sponges (*Porifera*). We found that the processes governing vertical transmission were both neutral and selective. A neutral model explained 66% of the variance in larval microbiota, which was higher than the variance this model explained for adult sponge microbiota (*R*^2^ = 27%). However, microbes that are enriched above neutral expectations in adults were disproportionately transferred to offspring. Patterns of vertical transmission were, however, incomplete: larval sponges shared, on average, 44.8% of microbes with their parents, which was not higher than the fraction they shared with nearby non-parental adults. Vertical transmission was also inconsistent across siblings, as larval sponges from the same parent only shared 17% of microbes. Finally, we found no evidence that vertically transmitted microbes are faithful to a single sponge host species. Surprisingly, larvae were just as likely to share vertically transmitted microbes with larvae from other sponge species as they were with their own species. Our study demonstrates that common predictions of vertical transmission that stem from species-poor systems are not necessarily true when scaling up to diverse and complex microbiomes.

## Introduction

All animals are colonized by microbes. These microbes live in communities, called microbiomes, that can exhibit astonishing diversity and complexity and have profound effects on host health and fitness [1, 2, 3]. However, despite their importance, we still do not understand how most organisms acquire their microbiomes: are they largely inherited from parents via vertical transmission, or acquired horizontally from the environment? In the last five years, the literature has provided widely divergent answers to this question [4, 5, 6, 7]. A recent meta-analysis of 528 host-microbe symbioses found that 42.8% of symbioses were strictly vertical, 21.2% were strictly horizontal, and 36% exhibited a combination of transmission modes [7]. Understanding how animals acquire their microbiomes, especially microbial symbionts, is necessary to learn how environments shape host phenotypes via host-microbe interactions and whether hosts and their microbiomes represent an important unit of natural selection [8, 9, 10, 11, 12].

Classic coevolutionary theory predicts that (i) if microbial symbionts are beneficial, they should be vertically transmitted (as the host is assured of gaining a compatible partner), and (ii) the more a host depends on its microbial partners, the higher the expected incidence of vertical transmission [13, 14, 15, 16, 17, 18, 19, 20]. In support, many obligate insect-microbe interactions, such as those described between *Buchnera-aphid* [21], *Wolbachia-nematode* [22], and *Ishikawaella-stinkbug* [23] are transmitted from parents to offspring. However, this theory is incomplete. Evidence for symbioses involving horizontal transmission is common, especially in hosts with relatively simple microbiota [6, 19, 24, 25, 26, 27]. Two well-known examples include the faculative symbiosis between the luminescent *Vibrio fischeri* and the bobtail squid *Euprymna scolopes* [28], and the obligate symbiosis between chemolithoautotrophic bacteria and the hydrothermal vent tubeworm *Riftia pachyptila* [29]. Furthermore, Mushegian and colleagues recently demonstrated that, in water fleas (*Daphnia magna*), microbes that are essential to host functioning are acquired from the environment and not maternally derived [30]. However, we currently do not understand whether the patterns and processes observed in these relatively species-poor systems can be extrapolated to highly diverse microbiomes. With increasing community diversity, do parents transmit a representative sample of the whole microbial community or select only a subset of the most beneficial microbes? How does vertical transmission interact with other community assembly processes shown to be important in complex communities, including ecological drift, priority effects, and environmental selection?

The present study is, to our knowledge, the first in-depth analysis of the nature, strength and consistency of vertical transmission in multiple coexisting host species in the wild from an animal phylum with diverse and abundant microbiomes. By characterizing signatures of vertical transmission in multiple, related host species, we also test, for the first time, partner fidelity between vertically transmitted microbes and their hosts. Partner fidelity is predicted by the hologenome theory of evolution because if vertically transmitted microbes occur in multiple host species, this weakens the coherence of the unit of selection [11]. Here we test these ideas in marine sponges, an evolutionarily ancient phylum with a fossil record dating back over 600 million years [31]. Indeed, Porifera are the oldest metazoan group with known microbial symbioses [32]. Marine sponges are filter feeders with a simple body plan consisting of canals embedded in an extracellular matrix called the mesohyl. Within the mesohyl, sponges maintain diverse microbial communities that contribute to host functioning by e.g., cycling nitrogen, fixing carbon dioxide, producing secondary metabolites, and acquiring and converting dissolved organic matter–tasks that, in many cases, the sponge cannot perform without microbial symbionts [32, 33, 34]. Sponge larvae are lecithotrophic, which means that they do not receive any external energy sources until they start filter feeding after metamorphosis and settlement [35]. However, some larvae travel long distances, much farther than what is expected from the energy content in their yolk. Evidence suggests that this is because some larvae have the capacity to gain energy via phagocytosis of vertically transmitted symbionts [36, 37, 38].

While the prevailing transmission model in marine sponges is a mixture of horizontal and vertical transmission [39], at least three lines of evidence suggest that vertical transmission could play an important role in the assembly of the sponge microbiome. First, sponges appear to have coevolved with a unique set of microbial symbionts that form so-called *sponge-enriched* 16S rRNA gene sequence *clusters* [40, 41]. These *sponge-enriched clusters* span 14 known bacterial and archaeal phyla, many of which are highly specific to the phylum Porifera (e.g., phyla such as Poribacteria, Chloroflexi and PAUC34f) [40, 41]. Unlike any other group of animal associated microbial symbionts described to date, each *sponge-enriched cluster* is monophyletic, indicating that microbes assigning to these clusters have diverged from their free-living relatives [40, 41]. Second, electron micrographs have revealed that sponge oocytes, embryos, and larvae contain free-swimming or vacuole-enclosed endosymbotic bacteria that are morphologically identical to those found in the mesohyl of the parent [42, 43, 44, 45, 37, 46]. The mechanisms of microbial selection and transference to oocytes vary between sponge species [46], as does the density and diversity of microbes that are incorporated into the oocytes [36, 47, 48]. Third, multiple studies, largely based on non-high-throughput sequencing methods, have found similar microbial phylotypes in adults and larvae from the same species [39, 49, 50, 51, 52]. One study also found that three pre-selected bacterial taxa that were present in the embryos of the tropical sponge *Corticium sp*. persisted throughout development and were consistently detected in adult samples over a period of three years [53]. Together, these lines of evidence strongly suggest that vertical transmission may be a frequent phenomenon that ensures the assembly of a functioning and beneficial microbiota in many species of marine sponges.

Here we use high-throughput sequencing to test for evidence of vertical and horizontal transmission by comparing microbial sharing in known parent-offspring pairs from wild sponges. We use these data to test four hypotheses of diverse microbiome acquisition that are applicable to any host-microbe system. Firstly, we test whether the processes underlying vertical and horizontal transmission are neutral or selective. If the processes underlying vertical transmission are neutral, microbes that are abundant inside the adults are expected to be widespread among their offspring. Alternatively, if the processes underlying vertical transmission are selective, then microbes found inside larval offspring should occur more frequently than expected, given their abundances in adults. Secondly, we test the hypothesis that sponges exhibit comprehensive vertical transmission, such that microbiota in larval offspring are either a perfect replica or a substantial subset of the microbes found in their adult parents. Alternatively, vertical transmission might be incomplete or undetectable; if incomplete, larval offspring will share only a fraction of their microbes with their parents, but this proportion will be higher than the proportion of microbes they share with other adults of the same species. If vertical transmission is undetectable, then larval offspring will be just as likely to share microbes with other conspecific adults as they are with their parents. Thirdly, we test the consistency of vertical transmission between parents and offspring. We hypothesize that if a specific set of symbionts has coevolved with their sponge host, and if it is adaptive for parents to transmit this set of symbionts, then all offspring from the same parent should receive an identical or highly consistent set of beneficial symbionts. Alternatively, if consistent vertical transmission is not important to parental fitness, or if parents benefit from transmitting different symbionts to each offspring (e.g., if larvae settle in variable environments where only a subset of symbionts is beneficial), then larvae might receive a variable or even random subset of microbes from their parents that is inconsistent between siblings. Finally, we test whether vertically transmitted taxa exhibit partner fidelity. If symbionts have coevolved with a particular sponge species, then conspecific sponge adults and larvae should share more vertically transmitted microbes with each other than with heterospecific hosts. Our approach helps reveal the prevalence and role of both vertical and horizontal transmission in an animal phylum with diverse microbiota that has important ramifications for understanding coevolution between hosts and their associated microbiota in general

## Results and Discussion

### Taxonomic diversity is distributed along a *sponge-specific axis*

To establish parent-offspring relationships for wild sponges, we placed mesh traps around adult sponges living close to the Islas Medas marine reserve in the northwestern Mediterranean Sea. We sampled 24 adults from a total of eight sponge species spanning five orders (Figure S1) and collected 63 larval offspring from 21 of these adults (1 to 5 larvae sampled per adult). To sample environmental microbes as a potential source pool, we simultaneously collected seawater samples from seven locations within the area where the adult sponges were found (seawater samples were never taken in direct proximity to any sponge specimen).

After quality control, we obtained 11,375,431 16S rRNA gene amplicon reads from these 94 samples (mean=121,015 reads per sample; min=1116, max=668,100 reads), resulting in 12,894 microbial ASVs (Amplicon Sequence Variants). Of these, 9,030 ASVs were present in the 24 sponge adults (Table S1), 5,786 were found in their 63 larval offspring (Table S2), and 9,802 ASVs occurred in the seven seawater samples. The 12,894 ASVs were classified to over 30 bacterial phyla and candidate phyla, five of which were only detected in the surrounding seawater. One class of Pro-teobacteria was unique to sponge adults, and two phyla, Deferribacteres and Fibrobacteres, were especially enriched in larval offspring, but present in low abundances in the other two environments (Figure 1A). While several phyla (classes for Proteobacteria) were shared between all three environments (circles close to the center in Figure 1A), likely representing horizontally acquired ASVs, a large fraction of the observed taxonomic diversity was only shared between sponge adults and larvae, distributed along a *sponge-specific axis* (left-hand side of the ternary plot in Figure 1A). These included many common sponge-associated phyla, such as Poribacteria, Chloroflexi, and PAUC34f, but also more arcane phyla like Tectomicrobia and SBR1093 (Figure 1A). Many of the sponge-associated phyla include microbes with known symbiotic features and functional capabilities. For example, members of Poribacteria and Chloroflexi harbor eukaryote-like protein domains which are suspected to be involved in preventing phagocytosis by the sponge host [54, 55]. Several genomic features in Chloroflexi are related to energy and carbon converting pathways, including amino and fatty acid metabolism and respiration, that directly benefit the sponge host [55]. Microbes from PAUC34f have the capacity to produce, transport and store polyphosphate granules, likely representing a phosphate reservoir for the sponge host in periods of deprivation [56]. This type of evidence strongly suggests that microbes from these phyla indeed represent beneficial symbionts for sponge hosts.

**Figure 1:**
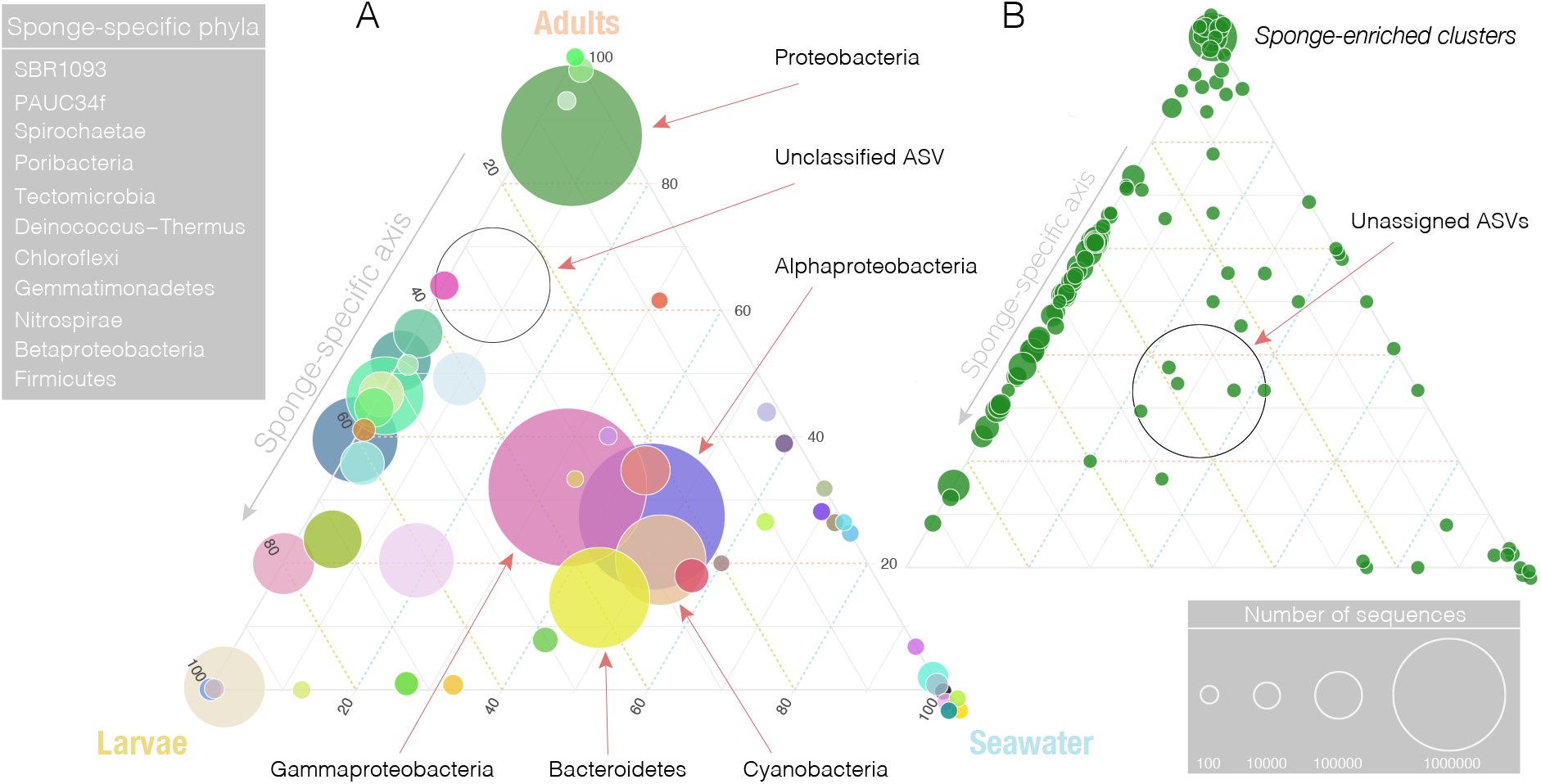
Ternary plots indicating the fraction of microbial ASVs classifying to (A) phyla, and assigning to (B) *sponge-enriched clusters*, present in three environments: seawater (light blue, bottom right corner); sponge adults (peach, top corner); and larval offspring (yellow, bottom left corner). Figure A shows the distribution of all microbial ASVs at the phylum level (class level for Proteobacteria). Each circle represents a different phylum, and the size of the circle corresponds to the total number of reads assigned to that phylum. While The color legend for (A) is shown in Figure 4. The phyla that lie along the sponge specific axis are listed in the grey table to the left of plot A. Figure B shows the diversity of all ASVs assigning to *sponge-enriched clusters*. Each green circle represents a different *sponge-enriched cluster*, and the size of the circle corresponds to the number of reads assigning to that particular cluster. ASVs that classify to phyla and *sponge-enriched clusters* that are unique to any of the three environments occur in their respective corners (100%); ASVs that classify to phyla and *sponge-enriched clusters* that are shared between any two environments occur along their focal axis. ASVs that classify to phyla and *sponge-enriched clusters* present in all three environments occur in the center of the ternary plots.

The ASVs we found also assigned to 105 different *sponge-enriched clusters* from 13 different bacterial phyla, of which Proteobacteria, Chloroflexi and Poribacteria represented the three most common (PAUC34f came in 5th place) (Figure 1B). These *sponge-enriched clusters* accounted for 9.6% of the total ASV richness and 25.5% of the total sequence count across samples. 94 *sponge-enriched clusters* were found in seawater, however, these only accounted for about 5% of ASV richness and 0.23% of reads from seawater. Out of these 94 *sponge-enriched clusters*, only 4 were not detected in the sponge hosts, supporting the idea that a rare biosphere functions as a seed bank for colonization of sponge hosts [57, 58]. While very few *sponge-enriched clusters* were present in all three environments (circles close to the center in Figure 1B), 62 were distributed along the *sponge-specific axis* (with a relative abundance of < 0.01% in the seawater). Sponge larvae do not filter feed prior to settlement and metamorphosis [35]. Concurrently, just one phyla and two *sponge-enriched clusters*, were shared between larvae and seawater only (bottom axis of Figure 1B), showing that, at least at these higher taxonomic levels, there is a signature of microbial transmission and subsequent enrichment between adults and larvae.

### The processes underlying symbiont acquisition are both neutral and selective

To test whether the processes underlying horizontal and vertical transmission are neutral or selective, we fit the neutral model developed by Sloan and colleagues [59] to adult and larval microbiota. This model predicts the relationship between the occurrence frequency of microbes in individual hosts (here either adults or larvae), and their abundances in a larger metacommunity consisting of microbes found in either (i) adults, including microbes that are shared with larvae and seawater (adults ∩ {larvae, seawater}; Figure 2A), or (ii) larvae, including microbes shared with adults and seawater (larvae ∩ {adults, seawater}; Figure 2B). Given neutral assembly processes, the model predicts that microbes with high abundances in the metacommunity should be frequently found in individual hosts. Microbes that fall above the neutral prediction occur more frequently than expected, indicating that they are selectively acquired and/or enriched by the sponge host, whereas microbes that fall below the neutral prediction occur less frequently than expected, and may therefore either be selected against or dispersal limited [60].

**Figure 2:**
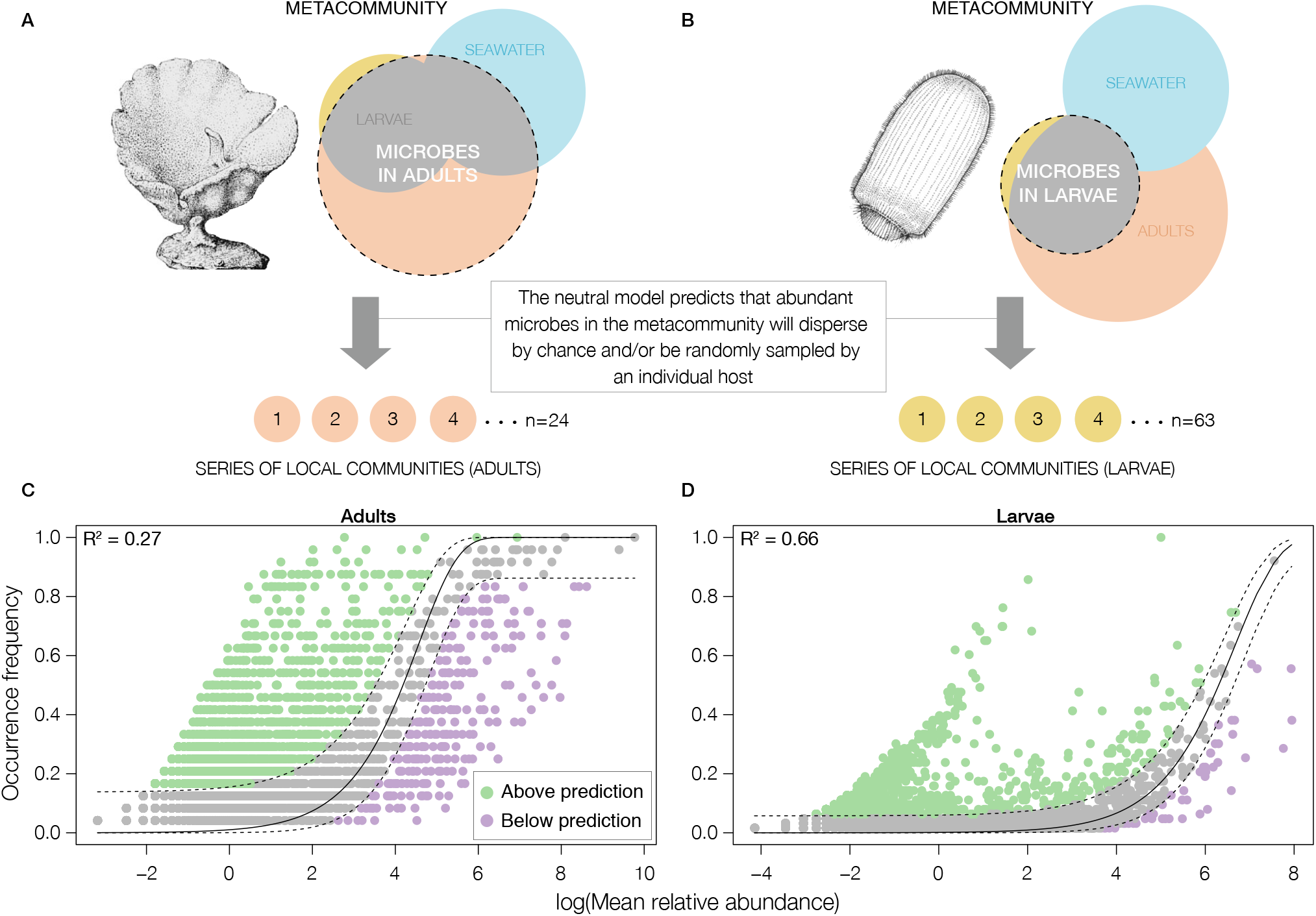
Panels A and B illustrate conceptual diagrams of the constructed metacommunities for (A) adults and (B) larvae. In the Venn diagrams, the microbial communities associated with adults, larvae, and seawater are depicted by circles colored in peach, yellow, and turquoise, respectively. The focal metacommunity is circled by a dashed black line, and the local host communities are represented as four circles below each Venn diagram, representing either individual adults (peach) or larvae (yellow). The bottom panel shows the fit of the neutral model for adults (C) and larvae (D). ASVs that fit the neutral model are colored gray; ASVs that occur more frequently than predicted by the model are colored green; and ASVs that occur less frequently than predicted are colored purple. Dashed black lines represent the 95% confidence interval around the model prediction (solid black line). The *R*^2^ for the model fit is shown in the upper left-hand corner of each plot. The percentage of microbes that fall above, within, and below the neutral prediction for adults and larvae are 23.7%, 73.4%, 3%, and 20.4%, 78.8%, 0.8%, respectively. This indicates that both neutral and non-neutral processes governs microbial acquisition in marine sponges.

In support of the hypothesis that neutral processes play an important role in vertical transmission in marine sponges, we found that the neutral model was a better fit to larval than to the adult microbiota (*R*^2^ = 0.27 in adults vs 0.66 in larvae; Figure 2C and Figure 2D). This pattern suggests that the importance of non-neutral processes increases as the sponge host matures, as mediated by selective acquisition of symbionts, active curation of the microbiota, and microbe-microbe interactions within the host. Evidence suggests that sponge hosts have mechanisms to actively recognise and incorporate symbionts into the oocytes [37, 61], but our results suggest that these mechanisms can be neutral and/or selective. Indeed, electron micrographs suggest that if microbes are collected by amoeboid nurse cells and subsequently engulfed by the oocytes, then the process may be selective [46, 62, 63, 61]. However, in the absence of nurse cells, microbes are incorporated into the oocytes solely based on their abundance [45, 64, 65, 66].

If the processes underlying vertical transmission are neutral, then microbes that are abundant in individual adults should be widespread among their larval offspring. To test this prediction, we examined whether microbes that fell above, within, and below the neutral prediction in adult sponges also were found within the same partitions across larvae (transmission of microbes goes from parent to offspring, not vice versa). We again found evidence that vertical transmission is governed by both neutral and selective processes. Owing to their filter feeding activities, adults harbor a large number of transient visitors, including food microbes (i.e., microbes that can be used as a food source by the host). Congruently, we found that 73.4% of the adult microbiota consisted of “neutral” ASVs (i.e., gray dots in Figure 2C). However, larvae only shared 41.7% of these ASVs; 10.7% and 88.5% fell above and within the neutral prediction. This suggests that microbes that are neutral in adults also tend to be neutral in larvae. In support of selective processes, of the 23.7% of ASVs that fell above the neutral prediction in adults (i.e., green dots in Figure 2C), 79% were present in larvae. Of these, 42.6% and 56.5% fell above and within the neutral prediction. While this indicates that symbionts that are selectively acquired and/or enriched by individual adults are also frequent across individual larvae, it also suggests that almost 43% are transmitted and incorporated into the oocytes by selective processes. The 10.7% that fell above the neutral prediction in larvae may represent symbionts that were haphazardly filtered by a few individual adults, and subsequently incorporated into the oocytes. Finally, for the 3% of ASVs that fell below the neutral prediction in adults (i.e., purple dots Figure 2C), 86.5% were present in larvae; 54.1% and 43.3% fell above and within the neutral prediction, suggesting that most of microbes that fall below the neutral prediction in adults represent dispersal limited symbionts.

From Figure S2A, an interesting pattern emerged: a “peak” consisting of microbes above the neutral prediction that largely disappeared when microbes shared with the seawater were removed (Figure S2B). Compared to adults, larvae do not filter feed prior to settlement and metamorphosis [35]. This, therefore, suggests that the “peak” consists of (i) a mixture of symbionts and other microbes that the adults acquire from the seawater (which are subsequently incorporated into the oocytes), and/or (ii) environmental microbes that populate the outer surface of the free-swimming larvae prior to settlement. While we can not exclude the latter, it is less likely for four reasons: (1) we rinsed sponge larvae with filter-sterilized seawater prior to DNA extraction; (2) evidence from electron micrographs suggests that microbes are not frequently present on the surface of sponge larvae [36, 44, 67]; (3) most of the ASVs forming the “peak” are also present above the neutral prediction in adults, indicating that they are selectively acquired and/or enriched by individual adults (Figure S3); and (4) several of these ASVs assigned to *sponge-enriched clusters* (Figure S4).

Our results allow to distinguishing between *direct* and *indirect* vertical transmission; that is, symbionts which have been passed down through multiple host generations, and microbes that the adult parent, at some point, acquired from the environment and subsequently incorporated into the oocytes. This parallels *direct* and *indirect* transmission in disease ecology [68], where a directly transmitted pathogen, e.g., HIV, moves directly from one host to another, without passing through the environment. An indirectly transmitted pathogen, such as *E. coli*, either requires time, or can survive indefinitely in the environment, before infecting a new host.

Furthermore, in systems with highly diverse microbiota, microbes are subject to continual turnover. This means that even vertically transmitted microbial lineages can be lost due to intrinsic and extrinsic factors affecting their population dynamics. Thus,*indirect* vertical transmission may provide a mechanism to either replenish lost or gain new symbiotic functions. With *directly* and *indirectly* vertically transmitted symbionts, offspring may also receive a larger microbial genetic repertoire. Similar to the idea that a genetically diverse cohort of offspring is more likely to succeed than a genetically uniform cohort, these larvae may better survive and reach adulthood in diverse and varying environments. However, to better understand *direct* and *indirect* vertical transmission, it will be necessary to trace transmission at the microbial strain level through multiple lineages of sponge hosts.

We next focused on the subset of vertically transmitted ASVs that were shared between adults and their offspring and were not detected in seawater (i.e., were least likely to be attributed to microbes from seawater found incidentally on the surface of the larva; Table S2). We found that 50.0%, 44.1% and 5.9% of these vertically transmitted ASVs fell above, within, and below the neutral prediction across individual adults, suggesting that at least half of the vertically transmitted ASVs are selected and/or enriched by individual adults. Of these “selected” taxa, 48.4% and 51.1% fell above and within the neutral prediction in larvae (Figure 3A and Figure 3B). Interestingly, of the ASVs that fell above the neutral prediction in adults, 40.6% assigned to *sponge-enriched clusters*, and of these, 51.7% and 48.3% fell above and within the neutral prediction in larvae (Figure 3 C and Figure 3D), further indicating that adults transmit beneficial symbionts to offspring that may likely be important during microbiome assembly. Of the 44.1% of ASVs that were neutral across individual adults, 22.3% and 77.2% fell above and within the neutral prediction in larvae (Figure 3A and Figure 3B), suggesting that offspring also receive “neutral” microbes that likely serve as an additional energy reserve until larval settlement. Finally, of the 5.9% of ASVs that fell below the prediction across individual adults, 42.3% and 53.8% fell above and within the neutral prediction in larvae (Figure 3A and Figure 3B). These percentages are altogether very similar to the ones found for the overall microbiota. This indicates that the relative importance of the neutral and selective processes that governs vertical transmission is similar regardless of whether microbes detected in seawater are considered or not.

**Figure 3:**
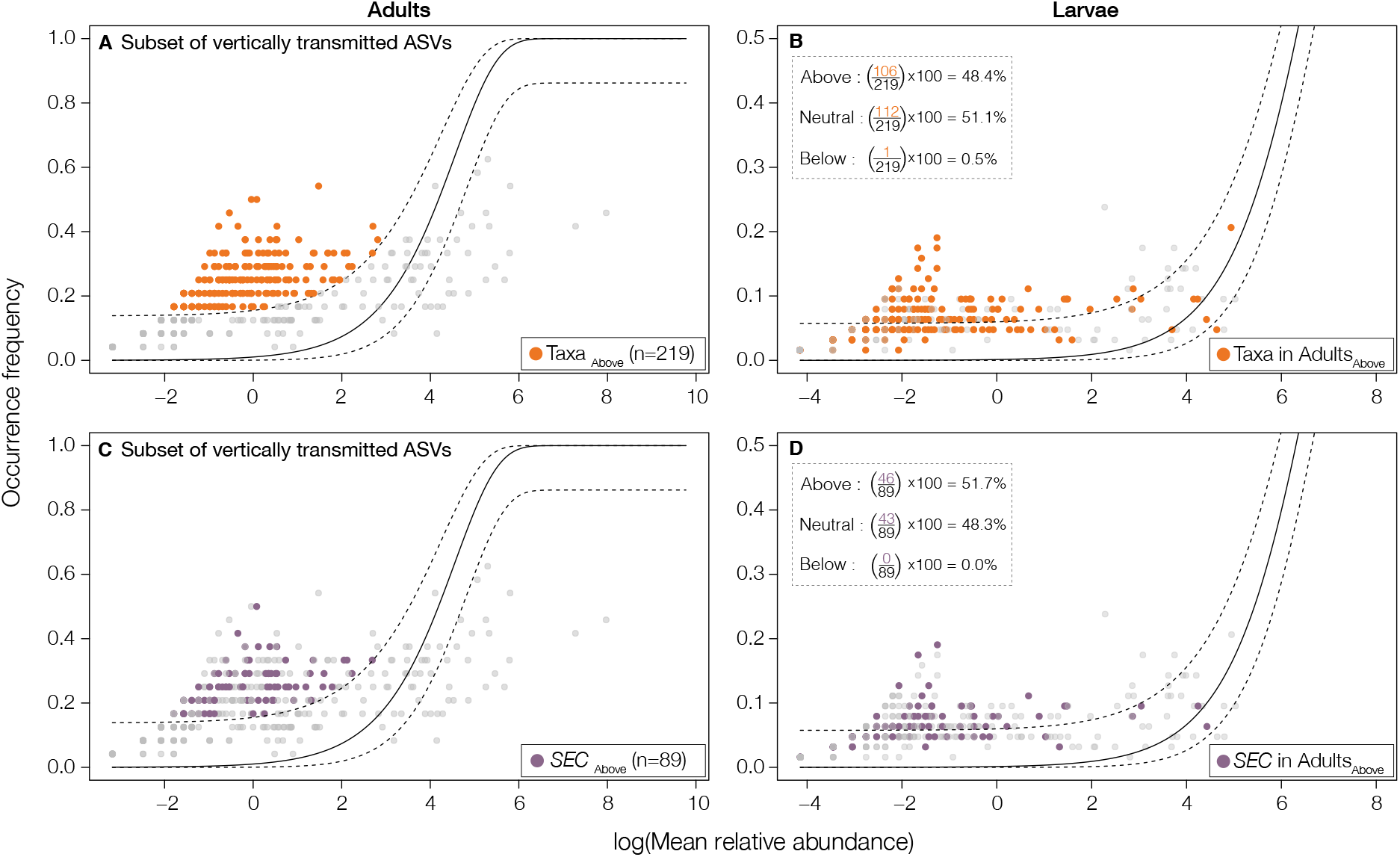
The processes underlying vertical transmission are both neutral and selective. Both the top and bottom panel correspond to ASVs shared between parents and their offspring, but not detected in seawater. In the top panel (A-B), orange dots correspond to ASVs that are above the neutral prediction in adults; 48.4% of these vertically transmitted ASVs are also above the neutral prediction in larvae. In the bottom panel (C-D), purple dots correspond to *sponge-enriched clusters* that are above the neutral prediction in adults; 51.7% of these vertically transmitted *sponge-enriched clusters* are also above the prediction in the larvae. Microbes that fall within or below the neutral prediction are colored in gray.

To further disentangle some of the processes underlying vertical transmission in marine sponges, in the remaining series of analyses, we introduce one broad (*overall*) and one narrow (*sponge-specific*) definition of vertical transmission (Figure S5A). In (1) *overall* vertical transmission, we consider all ASVs that are shared between parents and offspring, regardless of their presence in seawater (Figure S6 and Figure S7; Table S2). In (2) *sponge-specific* vertical transmission, we only include ASVs that are shared between parents and offspring, but were not detected in seawater (Figure S6 and Figure S7; Table S2). *Sponge-specific* vertically transmitted (VT) microbes are nested within the set of *overall* VT microbes. Specifically, *sponge-specific* VT microbes are restricted to symbionts that are not detected (or under detection limit) in seawater, including members of the rare biosphere, and *directly* VT symbionts. *Overall* VT microbes include transient microbes passing through the adult host, and symbionts which are selectively acquired from the seawater.

### Vertical transmission in sponges is relatively comprehensive, but often undetectable

We next tested whether patterns of vertical transmission were detectable in sponges, and if so, whether these patterns were comprehensive or incomplete (Figure S5B). A visual inspection of taxonomic profiles of the microbiota between parents and offspring indicated that offspring often harbor similar microbial phyla to their parents, as well as to non-parental conspecific adults (Figure 4 A and Figure S8). Adults and larvae were also fairly similar at the level of individual ASVs: across all sponge species, larvae shared, on average, 44.8% of their *overall* ASVs with their adult parents (Figure 5A and Figure S9). Parents and offspring also shared, on average, 60.7% of their *overall sponge-enriched clusters* (Figure S10A and Figure S11). These results suggest that vertical transmission is relatively comprehensive, at least when considering microbes also found in seawater. However, the percent of ASVs shared between parents and offspring was not different than the percent of ASVs and/or *sponge-enriched clusters* that larvae shared with conspecific adults living nearby (ASVs: 44.8% vs 44.5%, Δ=-0.34, 95% CI [−4.49,3.87], Mann-Whitney U=3917.5, P>0.1, Figure 5 A; *sponge-enriched clusters:* 60.7% vs 61.3%Δ=−0.36, 95% CI [−6.94,5.73], Mann-Whitney U=3916.5, P>0.1, Figure S10A). This pattern indicates that, at the level of all the microbes found in larvae, the signature of vertical transmission is essentially undetectable.

**Figure 4:**
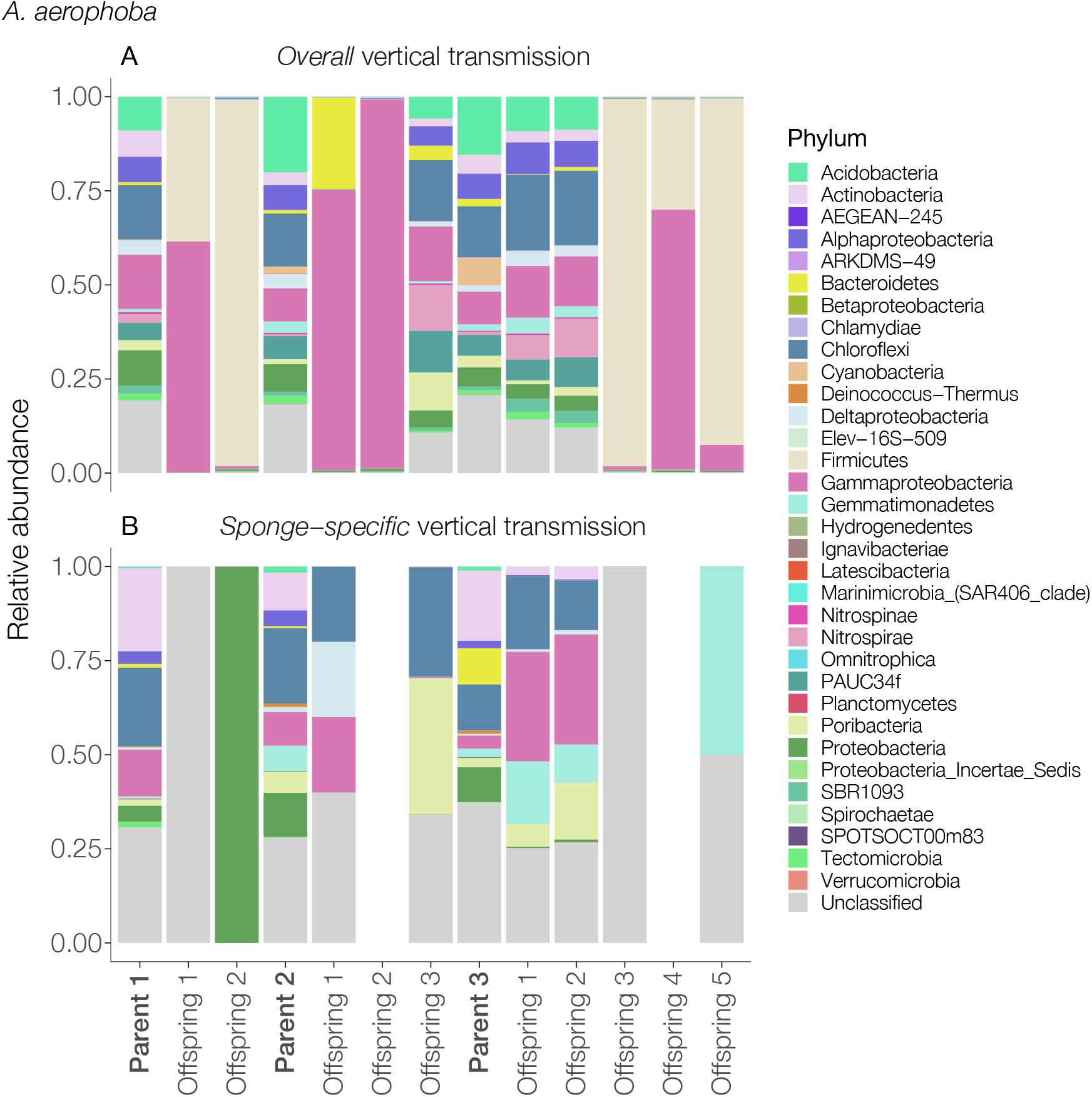
The relative contribution of vertically transmitted ASVs classifying to different microbial phyla (classes for Proteobacteria) in parents and the offspring of sponge species *A. aerophoba*. The top panel (A) shows the relative contribution of phyla for the *overall* definition of vertical transmission, and the bottom panel (B) shows the relative contribution of phyla for *sponge-specific* vertical transmission. Parents (in bold) and offspring are shown on the x-axis. Note that when microbes detected in seawater are removed, this sometimes leaves no vertical transmitted ASVs for the *sponge-specific* vertical transmission. Colors represent different microbial phyla (classes for Proteobacteria).

However, the analysis above included ASVs found in seawater, which may represent transient microbes passing through adult hosts, which are not consistent or important members of the sponge microbiota, or incidental microbes found on the surface of the larva. Removing ASVs detected in seawater not only reduced the taxonomic diversity found in larvae, but also decreased the percent of microbes shared between adults and their larval offspring. On average, offspring only shared 11.3% and 18.6% of their *sponge-specific* ASVs (Figure 5B and Figure S9) and *sponge-enriched clusters* (Figure S10B and Figure S11) with their parents. This pattern indicates that, for *sponge-specific* ASVs and *sponge-enriched clusters*, vertical transmission is incomplete. However, the detectability of vertical transmission in creased somewhat in these subsets. Specifically, the percent of *sponge-specific* ASVs shared between parents and offspring was slightly but significantly higher than the percent of *sponge-specific* ASVs larvae shared with nearby conspecific adults (11.3% vs 8.8%, Δ=2.23, 95% CI [0.00,5.00], Mann-Whitney U=4685, P=0.04; Figure 5B). However, parents and offspring did not share significantly more *sponge-specific sponge-enriched clusters* than with nearby conspecifics (18.6% vs 14.12%, Δ=0.00, 95% CI [-0.00,0.00]; Mann-Whitney U=4388, P>0.1; Figure S10B). These patterns persisted when we filtered the data to only contain samples with ≥5,000 reads (Figure S14). However, due to the large reduction in sample sizes, no significant differences were found in data filtered to only contain samples with ≥10,000 reads (Figure S15). When ASVs were agglomerated to the family or genus level, the results were similar for the *overall* vertical transmission (Figure S16A and Figure S16B). However, for the *sponge-specific* vertical transmission, the percents shared were lower than for taxa at the ASVs level (Figure S16A and Figure S16B). This is because many genera, and many more families, are shared with seawater, and the constraints implied by the *sponge-specific* definition of vertical transmission necessarily removes many of these higher-level taxa from the analysis.

**Figure 5:**
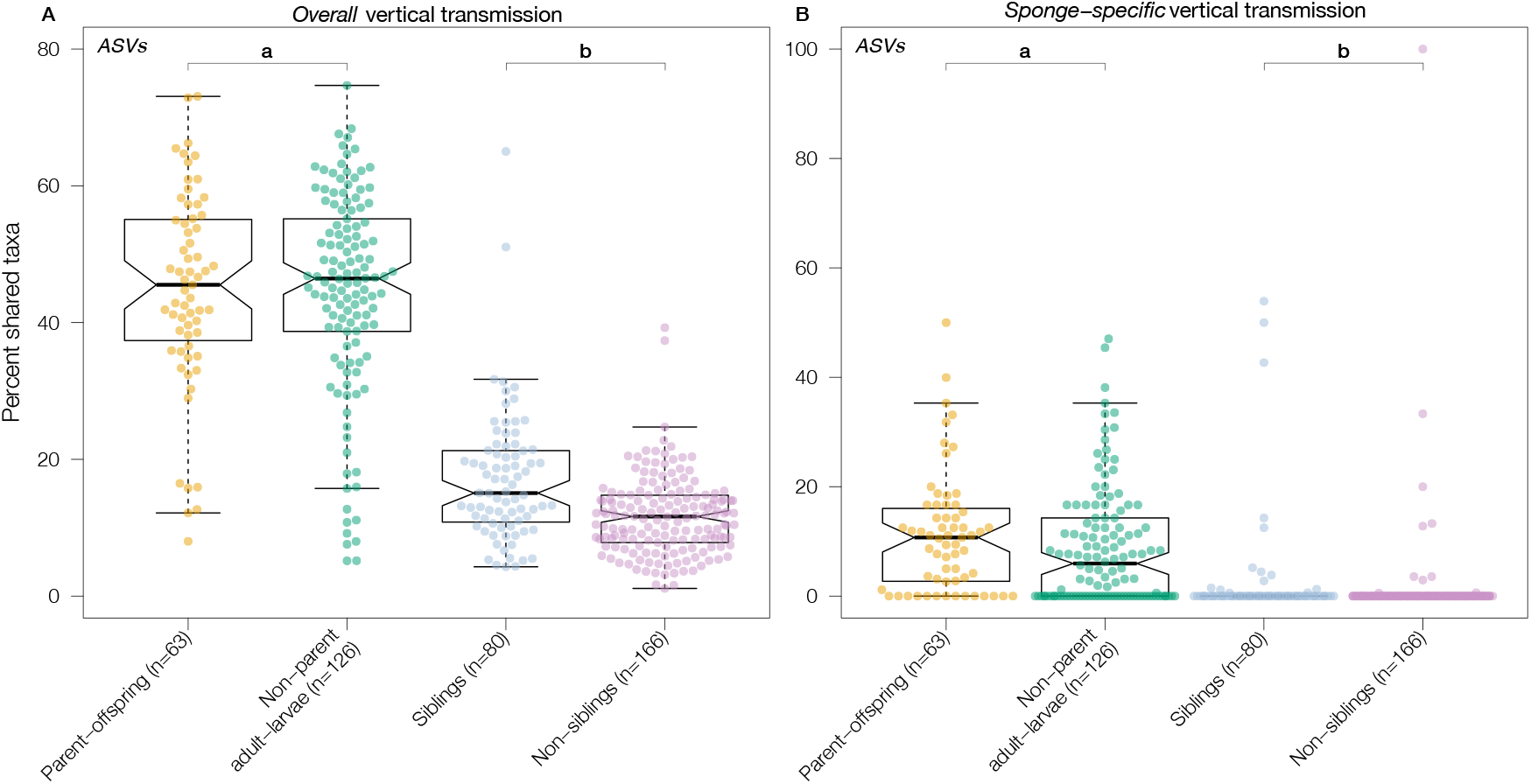
Percent of shared ASVs in the (A) *overall* and (B) *sponge-specific* definition of vertical transmission. Box-plots (a) show the percent shared ASVs between sponge larvae and either (i) their known parents (yellow dots), or (ii) non-parental conspecific adults (green dots). In boxplots (a), each dot represents one parent-offspring pair, or one non-parent adult-larva pair across all sponge species (see Figure S9). For *overall* vertical transmission (A), parents and offspring shared, on average, 44.8% of the ASVs, whereas non-parental conspecific adults and larvae shared, on average, 44.5% of ASVs (P>0.1). For *sponge-specific* vertical transmission (B), parents and offspring shared, on average, 11.3% of ASVs, whereas non-parental conspecific adults and larvae shared, on average, 8.8% of ASVs (P=0.04). Boxplots (b) show the percent shared VT ASVs between (i) siblings (blue dots), and (ii) non-siblings (purple dots). In boxplots (b), each dot represents one sibling pair, or one pair of non-siblings (see Figure S12). For *overall* vertical transmission (A), siblings shared, on average, 17.0% of their VT ASVs, while non-siblings only shared 11.7% (P<0.001). For *sponge-specific* vertical transmission (B), siblings shared, on average, only 2.4% of their VT ASVs, whereas non-siblings shared 1.0%(P=0.001). While these are significantly different, the effect size (i.e., the difference in location, Δ), is effectively zero.

To further characterize patterns of vertical transmission, we computed modularity on bipartite networks constructed for each sponge species. In the ecological network literature, modules are groups of species that “interact” more among themselves than with groups of other species (e.g., flowers and their pollinators, or fruits and their seed dispersers). If modules are perfectly separated; that is, no species interact with species from other modules, they are called compartments. Weighted modularity has been shown to be positively correlated with network specialization 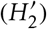, reinforcing the idea that modules exist because species only interact with a small number of other coevolved species [69]. Computing modularity on weighted bipartite networks allows for weighting species by their relative abundances, such that rare microbes are down-weighted and modules are formed around the most common host–microbe associations [69, 70]. We computed modularity on two sets of bipartite networks: (1) the *overall* networks which contain conspecific hosts (i.e., adults and larvae from the same species) and all ASVs detected in those hosts; (2) the *sponge-specific* networks that contain conspecific hosts and ASVs detected in those hosts, but not in seawater. If parents and offspring harbor the same set of microbes at similar abundances, and if those microbes are unique to a given set of parents and offspring, then the observed networks should be organized into compartments. We tested whether the observed modules deviated from these expected parent-offspring compartments using the Normalized Mutual Information (NMI) criterion [71, 72, 73]. NMI ranges between 0 and 1, where 0 indicates complete dissimilarity between expected compartments and observed modules, and 1 indicates that the observed modules only contain nodes corresponding to parents and offspring (i.e., compartments). We found that, while both the *overall* and *sponge-specific* networks were modular (*overall:* 0.48±0.17; *sponge-specific:* 0.57±0.14), the observed modules were not comprised of nodes corresponding to parents and offspring (Figure S17A). The *sponge-specific* networks had, on average, the highest NMI score but these networks were still quite far from the prior expectation of perfectly separated parent-offspring compartments (*overall*: 0.49±0.08 and *sponge-specific*: 0.36±0.13; Figure S18a). We also computed modularity on unweighted bipartite networks. While these resulted in different module composition, modules were still not comprised of nodes only corresponding to parents and offspring (Figure S17B and Figure S18b).

### Vertical transmission is largely inconsistent; but each offspring receives a small set of identical microbes from their parent

If symbiotic microbes have coevolved with their sponge host, and if it is adaptive for parents to transmit these microbes, then we would expect that all offspring from the same parent should receive an identical or highly consistent set of beneficial symbionts (Figure S5C). Alternatively, if consistent vertical transmission is not important to parental fitness, or if parents benefit from transmitting different symbionts to each offspring, then we would expect larvae to receive a variable or even random subset of microbes from their parents that is inconsistent between siblings.

We tested this prediction by calculating the proportion of *overall* and *sponge-specific* VT ASVs shared between larval siblings and conspecific non-sibling larvae. Across all sponge species, siblings shared, on average, 17.0% and 18.8% of their *overall* VT ASVs (Figure 5A and Figure S12A) and *sponge-enriched clusters* (Figure S10A and Figure S13A). These sharing percentages were much lower than what larvae shared with their parents, but they were significantly higher than the percents of VT ASVs and *sponge-enriched clusters* they shared with non-sibling conspecific larvae (ASVs: 17.0% vs 11.7%,Δ=4.48, 95% CI [2.68,6.26], Mann-Whitney U=9145, P=<0.001, Figure 5A; *sponge-enriched clusters*: 18.8% vs 12.4%, Δ=4.61, 95% CI [2.08,7.21], Mann-Whitney U=8531.5, P=<0.001, Figure S10A). This pattern indicates that, while each offspring receives a small number of identical microbes from their parent, VT microbes are largely inconsistent across siblings. When we removed ASVs detected in seawater, siblings and non-siblings only shared 2.4% and 1.0%, and 1.85% and 0.6% of their VT ASVs, respectively (Δ=0.00, 95% CI [-0.00,0.00], Mann-Whitney U=6383, P=0.001, Figure 5A and Figure S12A) and *sponge-enriched clusters* (Δ=0.00, 95% CI [-0.00,0.00], Mann-Whitney U=6076, P=0.024, Figure S10A and Figure S13A). Together, these results indicate that siblings receive a small set of identical symbionts, but that the majority of these microbes originate from the seawater where they have been selectively acquired by the adult parent prior to being transmitted to offspring.

The absence of a large consistent set of microbes transmitted between a given parent and its offspring could have at least three explanations. First, perhaps only a few symbiotic microbes are required to establish a functioning and beneficial microbiota; hence, parents might only “selectively” transmit a few of the most important symbionts to offspring. Second, parents may benefit from varying the microbes transmitted to each offspring. Such variability might be important if offspring disperse long distances and settle in diverse and varying environments. In this case, the identity of the most favorable set of microbes may vary across environments. This explanation is analogous to the idea that a genetically diverse cohort of offspring is more likely to succeed than a genetically uniform cohort (in this case, the genetic diversity is microbial, and not from the host). Third, previous research has suggested that larvae can phagocytose VT microbes [36, 38, 37]. Thus, to maximize their offspring’s chances of survival until settlement, parents may “neutrally” transmit a large number microbes as an additional energy source. All of these explanations are congruent with the finding that the mechanisms underlying vertical transmission are likely both neutral and selective.

### Vertically transmitted microbes are not host species-specific

By the time many sponges reach adulthood, they have converged on highly similar and species-specific microbiota [74, 75], including the eight sponge species analyzed here [74, 76, 77, 78]. These distinctive, species-specific communities may reflect the nature and strength of host-microbe interactions and strong selection for certain symbionts at the host species level. Furthermore, if this selection is a result of strong coevolution between microbes and hosts, then we would expect high levels of host species fidelity; that is, conspecific adults and larvae should share more VT microbes than they do with individuals from different host species (Figure S5D). While previous studies, largely based on non-high-throughput sequencing methods, have found similar microbial phylotypes in adults and larvae from the same sponge species [39, 49, 50, 51, 52], little is known whether this is also the case for larvae from multiple species.

We tested this prediction by calculating the percent of shared vertically transmitted ASVs among offspring from all possible combinations of adults. Surprisingly, we found that larvae were not more likely to share vertically transmitted ASVs (Figure 6A and Figure S19A) or *sponge-enriched clusters* (Figure S21A and Figure S22 A) with larvae from their own species as compared to larvae of other species (ASVs: 17.4% vs 15.5%, Δ=1.72,95% CI [-2.71,6.27], Mann-Whitney U=1928, P>0.1; *Sponge-enriched clusters:* 21.3% vs 18.6%, Δ=2.81, 95% CI [-2.93,8.67], Mann-Whitney U=1966.5, P>0.1). This pattern persisted when we considered the relative abundances of the vertically transmitted ASVs in larvae (6.6% vs 6.2%, Δ=-0.05, 95% CI [-1.07,1.04], Mann-Whitney U=1691, P>0.1, Figure 6A and Figure S19B)) and *sponge-enriched clusters* (20.4% vs 15.6%, Δ=3.75, 95% CI [-1.04,15.20], Mann-Whitney U=1973, P>0.1, Figure S21A and Figure S22B). Removing microbes detected in seawater, conspecific larvae shared, on average, only 3.5% and 9.5% of their *sponge-specific* vertically transmitted ASVs (Figure S20A) and *sponge-enriched clusters* (Figure S23A), respectively. These percents of sharing were not different than the percent of vertically transmitted ASVs larvae from different species shared (ASVs: 3.5% vs 2.7%, Δ=-0.000, 95% CI [-0.41,0.00], Mann-Whitney U=1651, P>0.1, Figure 6B; *sponge-enriched clusters:* 9.5% vs 8.6%, Δ=-0.000, 95% CI [-2.65,3.61], Mann-Whitney U=1696, P>0.1, Figure S21B). Similar results were also observed when we considered the relative abundance of vertically transmitted ASVs (5.0% vs 1.8%, Δ=-0.000, 95% CI [-0.01,0.00], Mann-Whitney U=1614.5, P>0.1, Figure 6B and Figure S20B) and *sponge-enriched clusters* (10.1% vs 8.5%, Δ=-0.000, 95% CI [-0.07,0.01], Mann-Whitney U=1591, P>0.1, Figure S21B and Figure S23B). These patterns persisted when we filtered the data to only samples with ≥ 5,000 and ≥ 10,000 reads (Figure S24 and Figure S25).

**Figure 6:**
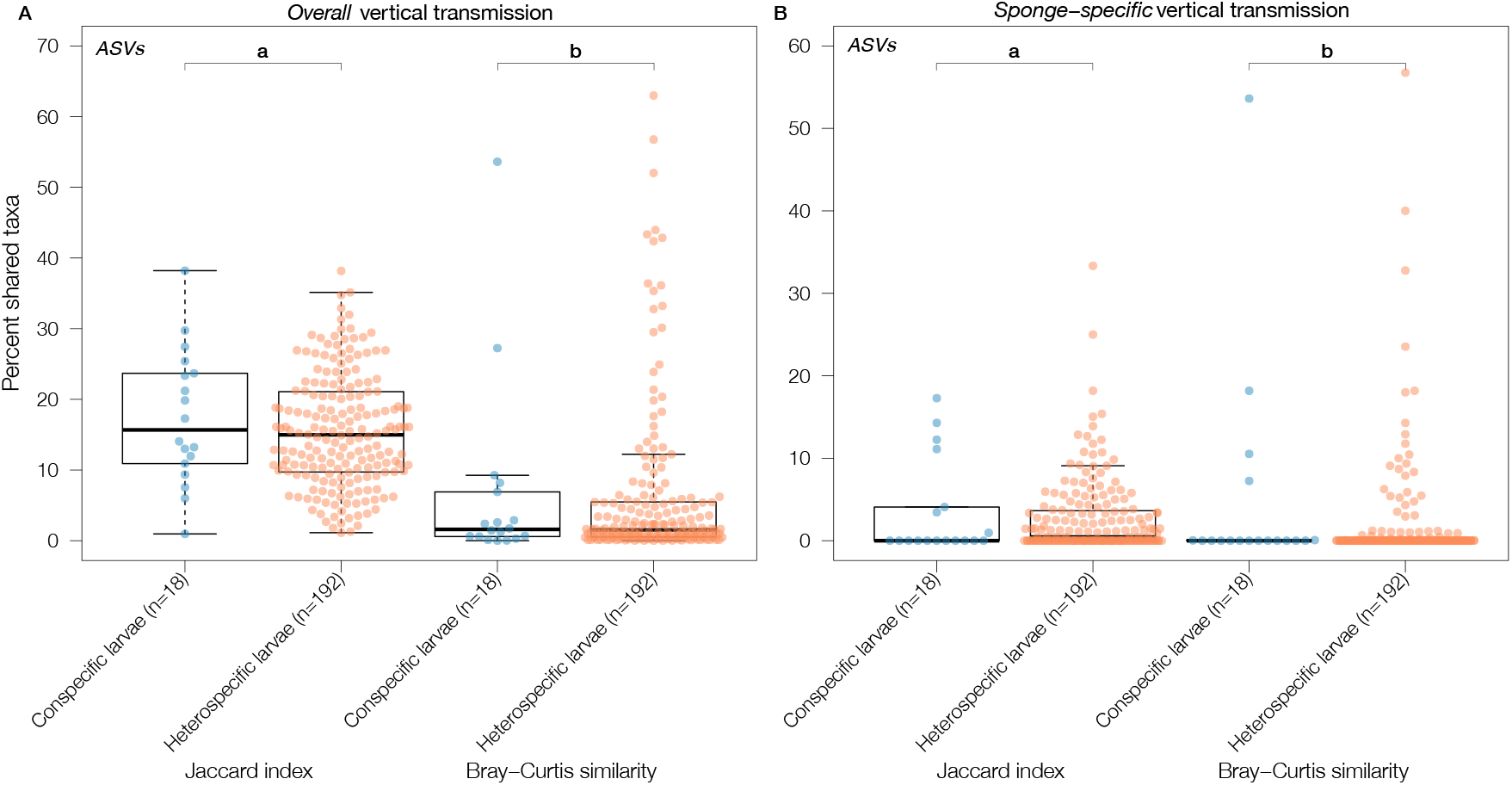
The percent of shared (A) *overall* and (B) *sponge-specific* vertically transmitted ASVs among offspring from all possible combinations of adults, calculated as either the (a) Jaccard index (see Figure S19A and Figure S20A), or (b) Bray-Curtis similarity (see Figure S19B and Figure S20B). Each dot represents all offspring from either (i) adults belonging to the same species (blue dots), or (ii) adults from different species (orange dots). While the Jaccard index calculates similarity between two samples based on the presence-absence of taxa, Bray-Curtis similarity weights taxa by their relative abundance. For *overall* vertical transmission (A), conspecific larvae shared, on average, 17.4% (Jaccard) and 6.6% (Bray-Curtis) of the ASVs, whereas heterospecific larvae shared, on average, 15.5% (Jaccard) and 6.2% (Bray-Curtis) of the ASVs (P>0.1). For *sponge-specific* vertical transmission (B), parents and offspring shared, on average, 3.5% (Jaccard) and 5.0% (Bray-Curtis) of the ASVs, whereas non-parental conspecific adults and larvae shared, on average, 2.7% (Jaccard) and 1.8% (Bray-Curtis) of the ASVs (P>0.1).

To test this pattern beyond pairwise comparisons, we computed weighted modularity on two bipartite networks: (i) the *overall* network which contains all hosts (i.e., adults and larvae) and ASVs detected in those hosts, and (ii) the *sponge-specific* network that contains all hosts and ASVs detected in those hosts, but not in seawater. If conspecific adults and larvae harbor the same microbes at similar abundances, and do not share those with other species, then the networks will be organized in compartments consisting of conspecific adults and larvae only. While we found that both the *overall* and *sponge-specific* network were highly modular (Q=0.71 and Q=0.79; Table S3), modules rarely consisted of adults and larvae from the same species (NMI=0.51 and NMI=0.41; Figure S26A). For instance, in the *overall* network, apart from *A. aerophoba’s* and *I. oros’s* adults that together formed one module, all other adults formed their own species-specific modules. In the *sponge-specific* network, there were a few modules that contained a larger number of heterospecific larvae (Figure S26A). In both networks, some modules that contained adults also contained larvae, but they rarely corresponded to offspring or even larvae of the same species. While the results from modularity computed on unweighted networks where quantitatively different (Table S3), it did not change the overall conclusion (Figure S26B).

## Conclusion

Vertical transmission is proposed to be a primary mechanism by which parents transmit assemblages of beneficial microbes to offspring in a way that maintains both these microbes’ interactions with each other and the beneficial functions that emerge from their interactions [16, 17, 19]. While this may often be the case when microbial symbionts consist of just one or a few species [21, 22, 23], when microbiomes are highly diverse, transferring hundreds to thousands of microbial species such that their interaction structures and emergent functions are preserved seems highly improbable. In systems with diverse microbiomes, how do parents ensure that offspring get the microbes they need? In marine sponges, we know that such mechanisms exist because by the time juveniles reach adulthood, they have converged on highly similar and species-specific microbiomes [74, 75]. Alternatively, species have evolved to create environments which select for specific microbes. Our results indicate that the assembly processes can be both neutral and selective, and that this may help to explain why several of our findings cast doubt on the consistency and faithfulness of vertical transmission of highly diverse microbiomes. Specifically, across eight sponge species, we show that: (1) vertical transmission is relatively comprehensive, but often undetectable. While larval sponges shared, on average, 44.8% of microbes with their parents, this fraction was not higher than the fraction they shared with nearby conspecific adults who were not their parents; (2) vertical transmission is inconsistent across siblings, as larval sponges from the same parent only shared 17% of microbes, and (3) vertically transmitted microbes are not faithful to a single sponge species: surprisingly, larvae were just as likely to share vertically transmitted microbes with larvae from other species as they were with their own species, and are therefore unlikely to have coevolved with particular sponge species. While removing microbes detected in seawater increased the detectability of vertical transmission, it heavily decreased the percent of microbes shared between adults and their larval offspring, including vertically transmitted microbes shared between siblings. Together, our results indicate that siblings receive a small set of identical symbionts, but that the majority of these microbes originate from the seawater where they likely have been selectively acquired by the adult parent prior to being vertically transmitted to offspring.

Our findings highlight the need for new theory that is specific to the acquisition and transmission of diverse microbiomes (see e.g., [79]), but also theory that not only considers horizontal and vertical transmission of microbes, but also *direct* and *indirect* vertical transmission; that is, symbionts which have been passed down through multiple host generations, and microbes that the adult parent, at some point, acquired from the environment and subsequently incorporated into the oocytes. While this mixture of mechanisms likely reduces the strength and consistency of vertical transmission, it may have other benefits that increase the chances of dispersing larvae to settle and reach adulthood in diverse and varying environments. For example, which microbes first colonize the host strongly influence subsequent community succession and stability [80, 81, 82, 83].

Finally, some of our results are relevant to the predictions put forward by of the hologenome theory of evolution [8, 9, 12]. This theory proposes that there might be value in treating hosts and their microbiota as a single evolutionary unit. This theory comes with an important expectation: high partner fidelity–if the collection of genomes varies within and between host generations, then it is not a coherent unit of selection [10, 11]. Such tight partner fidelity is typically only found among host-microbe symbioses with obligate vertical transmission. On the contrary, we found that many vertically transmitted microbes, including many *sponge-enriched clusters*, were not faithfully transmitted by parents to offspring nor were they host species-specific. As such, their evolution is likely shaped by multiple host species across the phylum Porifera, as well as by the marine environment where the sponge hosts live. It remains to be further tested whether the patterns reported here hold for even more sponge species, or persist when using larger sequencing depths or strain tracking techniques. Overall, our study demonstrates that common predictions of vertical transmission that stem from species-poor systems are not necessarily true when scaling up to diverse and complex microbiomes.

## Methods

We collected sponge and seawater samples between July and August 2012, close to the Islas Medas marine reserve in the northwestern Mediterranean Sea 42°3′0″*N*, 3°13′0″*E* by SCUBA at depths between 5-15 m. The analyzed species are common Mediterranean sponges and were identified based on their distinct morphological features. The sampling site consisted of a relatively small bay (roughly 18,000 *m*^2^). All sampled sponge species live in rocky overlapping habitats, and all species could be found within the same depth range. However, some specimens were found in more shaded areas than others.

### Larval sponge collection

We constructed larvae traps by modifying the traps used in [84] (Figure S27). In order to collect offspring from known parents, traps were mounted over individual adult sponges by SCUBA. To minimize stress to individual adults, traps were removed after one week. During this time, sample bottles were collected and replaced each day. Bottles were placed on ice in insulated coolers and transported to the laboratory (< 2 hours). Larvae were identified using a stereolupe. In order to remove loosely associated microbes, larvae were carefully rinsed with filter-sterilized seawater (0.20 *μ*m filter) before preservation in RNA later. All larval samples were stored at −80°C until DNA extraction.

### Adult sponge collection

After larval offspring were collected, three adults per sponge species were sampled. These individuals corresponded to the same adults from which we collected larvae. However, for a few species, larvae could only be collected for two adults. In these cases, a third adult was still sampled. Specimens were sub-lethally sampled by removing a small sample of tissue. Excised tissue was placed in separate plastic tubes and brought to the surface where they were preserved in RNA later and placed on ice in insulated coolers and transported to the laboratory (< 2 hours). Seawater samples were collected at 5 m depth and at seven locations within the sampling area. The water was always collected at deeper locations (> 5 m) within the sampling area, and never in direct proximity to the benthic community. All seven water samples were poured into separate, sterile 5 L jars. Aliquots of seawater (300-500 mL each, 1 aliquot per sample jar) were concentrated on 0.2 *μ*m polycarbonate filters, and submerged in lysis buffer. All samples were stored at −80^°^C until DNA extraction.

### DNA extraction and sequencing

DNA was extracted from ≈0.25 g of adult sponge tissue using the PowerSoil DNA extraction kit (MoBio). DNA from larvae (one larva per adult) was extracted using the XS-RNA extraction kit (Macherey-Nagel) because of its capacity to extract DNA from small samples, i.e., one larva. All DNA extractions were performed according to standard protocols. The seven seawater samples were processed by passing 2 L of seawater through 0.2 *μ*m Sterivex filters, and DNA was extracted from these filters as described by [52]. All extractions included a negative control without sponge tissue, and the lack of amplified DNA was examined with the universal bacterial primers 27F and 1492R. The V4 region of the 16S rRNA gene was amplified using the primer set 515FB-806RB [85], and sequenced using the Illumina HiSeq2500 platform. Sequencing was performed by the Earth Microbiome Project [86].

### Sequencing analysis

Illumina-sequenced, single-read fastq files were processed and cleaned in `R` [87] using the default settings in `DADA2` [88] to produce an amplicon sequence variant (ASV) table (Appendix 1), and `Silva` (v128) [89] was used to create the ASV taxonomy. The `Phyloseq R` package [90] was used to filter out sequences classifying to *Archaea* and *Eukaryota*. We also removed singleton ASVs, and phyla that occurred in less than two samples (Appendix 1). The analyzed dataset contained samples with at least 1,000 reads.

### Identification of *sponge-enriched clusters*

A representative sequence from each ASV was taxonomically assigned using a BLAST 62 search against a curated ARB-SILVA database containing 178 previously identified *sponge-specific clusters* [41]. For each BLAST search, the 10 best hits were aligned to determine sequence similarities. The most similar ASV sequence to the respective reference sequence within the database was then assigned to an *sponge-specific clusters* based on a 75% similarity threshold: (i) a sequence was only assigned to any given *sponge-specific clusters* if its similarity was higher to the members of the cluster than to sequences outside the cluster; and (ii) if its similarity to the most similar sequence within the cluster was above 75%. A majority rule was applied in cases where the assignment of the most similar sequences was inconsistent, and the ASV sequence was only assigned to the *sponge-specific clusters* if at least 60% of the reference sequences were affiliated with the cluster.

### Data analyses

We used `Phyloseq` package in `R` to store, organize and filter the analyzed sequence data [90]. Furthermore, to find ASVs corresponding to our two definitions of vertical transmission, we used set theory functions in `R`, e.g., `setdiff(A, B)` to find all features in `A` that is not present in `B` (i.e., `A \ B`), or `intersect(A, B)` to find all features that are present in both `A` and `B` (i.e., `A ∩ B`). We used box plots overlaid with swarm plots (using the `beeswarm` package in `R`) to better visualize the distribution of the data. When enough data points existed (or where the confidence interval (i.e. size of the notches) was not larger than the interquartile range), we used the `boxplot (…, notch=T)` to draw a notch in each side of the boxes. If the notches of two box plots do not overlap, this can be seen as “strong evidence” that the two medians differ. To accompany this, we perform the Mann-Whitney U test, `wilcox.test(x, y, …)`, and to further compute a nonparametric confidence interval (CI) and estimate the difference in location between parameters `x` and `y` (i.e., the effect size, Δ), we set the argument `conf.int=T`. Modularity was computed using the `DIRT_LPA_wb_plus` algorithm in `R` [70]. To test whether observed modules deviated from prior expectations, we used Normalized Mutual Information (NMI) criterion calculated through the `NMI::NMI(x, y)` function in `R`.

## Acknowledgements

We thank Dr. Rafel Coma and Dr. Eduard Serrano for help in the field. J.R.B. was supported by an FPI Fellowship from the Spanish Government (BES-2011-049043). J.M.M. was supported by the French LabEx TULIP (ANR-10-LABX-41; ANR-11-IDEX-002-02), by the Region Midi-Pyrenees project (CNRS 121090) and by the FRAGCLIM Consolidator Grant, funded by the European Research Council under the European Union’s Horizon 2020 research and innovation programme (grant agreement number 726176).

## Authors’ contributions

J.R.B. and J.M.M. conceived the study. J.R.B. performed the fieldwork and analyzed the data. J.R.B. and J.M.M. drafted the first versions of the manuscript, and J.R.B. and E.A. refined the ideas and wrote the final version of the paper. C.D. helped in the field and extracted DNA from the larvae. C.A.G. identified the *sponge-specific clusters*. All authors commented and approved of later versions of the paper.

## Data and code availability

All data and code will be available on Open Science Framework with an R Markdown document such that all analyses and figures can be reproduced.

