## Supplementary material for "Vertical transmission of sponge microbiota is inconsistent and unfaithful"

**This file includes:**

supplementary tables 1-3

supplementary figures 1-27

Table S1: The number of sequence reads, and the number ASVs for (a) all microbes detected in the focal adult, including those detected in seawater, and (b) microbes found in the focal adult but not detected in seawater. Each row corresponds to an adult sponge. The last set of rows correspond to the seven seawater samples.

| Sponge species | $N_{reads}$ | $N^{a}_{ASVs}$ | $N^{b}_{ASVs}$ |
|---|---|---|---|
| | 144148 | 1581 | 588 |
| *Aplysina aerophoba* | 194322 | 1539 | 668 |
| | 150522 | 1334 | 581 |
| | 283401 | 2153 | 906 |
| *Ircinia oros* | 287524 | 1981 | 910 |
| | 299987 | 2217 | 867 |
| | 89896 | 1027 | 356 |
| *Ircinia fasciculata* | 182657 | 1319 | 439 |
| | 256141 | 1557 | 563 |
| | 48893 | 1130 | 200 |
| *Crambe crambe* | 180038 | 1099 | 149 |
| | 190642 | 810 | 104 |
| | 176885 | 490 | 70 |
| *Cliona viridis* | 147933 | 654 | 103 |
| | 140974 | 748 | 86 |
| | 70402 | 1512 | 164 |
| *Dysidea avara* | 127357 | 1921 | 224 |
| | 4232 | 324 | 40 |
| | 536732 | 2247 | 337 |
| *Hemimycale columella* | 114731 | 879 | 114 |
| | 1797 | 270 | 34 |
| | 149832 | 1333 | 248 |
| *Oscarella lobularis* | 330103 | 1779 | 293 |
| | 521058 | 2558 | 489 |
| | 236409 | 3507 | |
| Seawater | 198260 | 4448 | |
| | 176447 | 5978 | |
| | 208076 | 4766 | |
| | 668100 | 3512 | |
| | 661187 | 3611 | |
| | 499139 | 3928 | |

Table S2: The number of sequence reads, and the number ASVs those reads assigned to for (1a) all microbes detected in the focal larva, including those shared with seawater, (1b) microbes found in the focal larva but not detected in seawater, (2a) all microbes shared between the focal larva and its parent, including those shared with seawater, and (2b) microbes shared between the focal larva and its parent but not detected in seawater. Note that no larvae were collected for adult 3 for species *I. oros* and *I. fasciculata*, and adult 2 for *H. columella*.

| Sponge species | Source adult | $N_{reads}$ | $N^{1a}_{ASVs}$ | $N^{1b}_{ASVs}$ | $N^{2a}_{ASVs}$ | $N^{2b}_{ASVs}$ |
|---|---|---|---|---|---|---|
| *A. aerophoba* | Adult 1 | 5227 | 68 | 9 | 28 | 1 |
| | | 1849 | 59 | 5 | 38 | 1 |
| | Adult 2 | 27846 | 206 | 35 | 118 | 5 |
| | | 3058 | 52 | 1 | 33 | 0 |
| | | 169456 | 954 | 198 | 473 | 63 |
| | Adult 3 | 439760 | 1456 | 480 | 577 | 159 |
| | | 267481 | 1337 | 438 | 560 | 175 |
| | | 18199 | 154 | 20 | 85 | 1 |
| | | 5801 | 78 | 6 | 37 | 0 |
| | | 29836 | 161 | 16 | 72 | 2 |
| *I. oros* | Adult 1 | 10397 | 85 | 8 | 55 | 1 |
| | | 11260 | 77 | 6 | 51 | 3 |
| | | 221306 | 790 | 125 | 318 | 23 |
| | | 17141 | 185 | 30 | 106 | 5 |
| | | 32123 | 211 | 28 | 116 | 3 |
| | Adult 2 | 55825 | 299 | 46 | 178 | 12 |
| | | 16737 | 82 | 7 | 50 | 1 |
| | | 15971 | 145 | 17 | 78 | 6 |
| *I. fasciculata* | Adult 1 | 6008 | 103 | 9 | 34 | 1 |
| | | 6899 | 87 | 12 | 44 | 2 |
| | Adult 2 | 5479 | 62 | 10 | 32 | 0 |

*Continued on next page*

Table S2 – *Continued from previous page*

| Sponge species | Source adult | $N_{reads}$ | $N^{1a}_{ASVs}$ | $N^{1b}_{ASVs}$ | $N^{2a}_{ASVs}$ | $N^{2b}_{ASVs}$ |
|---|---|---|---|---|---|---|
| *C. crambe* | Adult 1 | 5272 | 114 | 8 | 33 | 1 |
| | | 2709 | 45 | 6 | 15 | 1 |
| | | 17264 | 186 | 26 | 71 | 4 |
| | | 2976 | 134 | 7 | 61 | 1 |
| | Adult 2 | 3054 | 97 | 8 | 46 | 1 |
| | Adult 3 | 18918 | 220 | 32 | 79 | 1 |
| | | 54956 | 419 | 59 | 147 | 2 |
| *C. viridis* | Adult 1 | 88167 | 284 | 40 | 86 | 2 |
| | | 2083 | 126 | 12 | 54 | 0 |
| | | 6669 | 96 | 7 | 37 | 0 |
| | | 6631 | 113 | 12 | 46 | 0 |
| | Adult 2 | 193439 | 443 | 53 | 162 | 5 |
| | | 77960 | 315 | 55 | 102 | 2 |
| | | 2961 | 145 | 14 | 70 | 1 |
| | Adult 3 | 42482 | 129 | 12 | 54 | 1 |
| | | 3517 | 61 | 8 | 29 | 0 |
| | | 5187 | 90 | 14 | 42 | 0 |
| *D. avara* | Adult 1 | 12651 | 66 | 11 | 23 | 0 |
| | | 30050 | 244 | 36 | 101 | 1 |
| | | 8630 | 67 | 4 | 31 | 0 |
| | Adult 2 | 78829 | 307 | 25 | 179 | 7 |
| | | 49906 | 234 | 34 | 112 | 4 |
| | | 44314 | 235 | 19 | 128 | 2 |
| | | 16470 | 225 | 38 | 94 | 1 |

*Continued on next page*

Table S2 – *Continued from previous page*

| Sponge species | Source adult | $N_{reads}$ | $N^{1a}_{ASVs}$ | $N^{1b}_{ASVs}$ | $N^{2a}_{ASVs}$ | $N^{2b}_{ASVs}$ |
|---|---|---|---|---|---|---|
| | Adult 3 | 37901 | 229 | 24 | 29 | 1 |
| | | 9215 | 113 | 13 | 18 | 1 |
| | | 27264 | 228 | 21 | 36 | 0 |
| | | 6937 | 91 | 13 | 15 | 0 |
| | | 372660 | 1111 | 258 | 89 | 3 |
| *H. columella* | Adult 1 | 459628 | 2245 | 769 | 803 | 72 |
| | | 71813 | 316 | 48 | 176 | 9 |
| | | 20188 | 139 | 17 | 5 | 91 |
| | | 349906 | 1468 | 519 | 570 | 45 |
| | Adult 3 | 9300 | 74 | 22 | 9 | 0 |
| *O. lobularis* | Adult 1 | 1218 | 79 | 6 | 46 | 1 |
| | | 6691 | 78 | 11 | 34 | 3 |
| | Adult 2 | 2872 | 79 | 5 | 42 | 0 |
| | | 110912 | 537 | 64 | 265 | 12 |
| | Adult 3 | 437458 | 1798 | 629 | 764 | 75 |
| | | 1116 | 52 | 8 | 38 | 1 |
| | | 2948 | 59 | 9 | 43 | 1 |
| | | 26825 | 151 | 13 | 92 | 1 |

Table S3: Weighted and unweighted modularity ($Q$), and Normalized Mutual Information criteria (NMI) for networks corresponding to the *overall* (O) and *sponge-specific* (SS) definition of vertical transmission, respectively.

| | Network | Number of modules | $Q$ | NMI |
|---|---|---|---|---|
| Weighted | O | 30 | 0.71 | 0.51 |
| | SS | 26 | 0.79 | 0.42 |
| Unweighted | O | 5 | 0.33 | 0.20 |
| | SS | 6 | 0.39 | 0.20 |

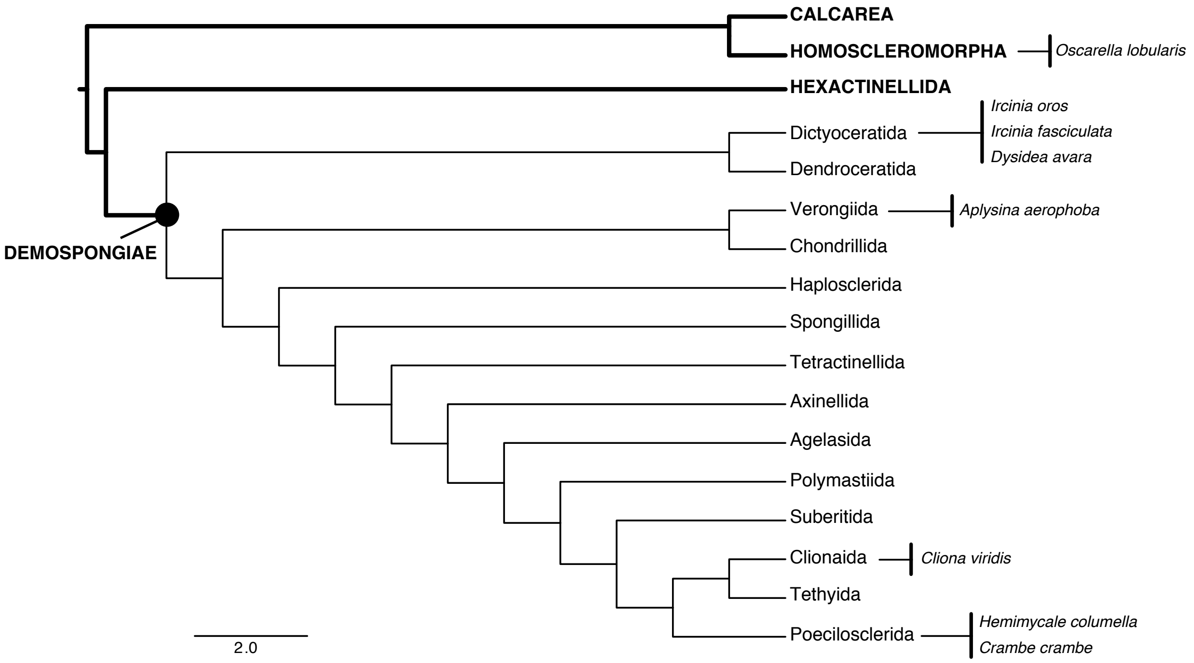


Figure S1: The phylogenetic relationship between the analyzed sponge species. The thin branches display orders within the class Demospongiae which contain over 75% of all sponges species worldwide. The eight analyzed sponge species span two classes (Homoscleromorpha and Demospongiae) and five orders (Homosclerophorida, Dictyoceratida, Verongiida, Clionaida and Poecilosclerida). All of the eight sponge species have the core of their known species range distribution within the Mediterranean Sea (which includes the Western Mediterranean Sea, the Adriatic Sea, the Ionian Sea, the Aegean Sea, and the Levantine Sea). More specifically, *Oscarella lobularis* has its known distribution in the Mediterranean Sea, part of the North Sea (the Swedish west coast), and part of the South Atlantic Ocean (the Azores, the Canary Islands, and Cape Verde); *Ircinia oros* has its known distribution in the Mediterranean Sea, and part of the south Atlantic Ocean (the Canary Islands); *Ircinia fasciculata* has its known distribution in the Mediterranean Sea; *Dysidea avara* has its known distribution in the Mediterranean Sea, the Black Sea, and part of the South Atlantic Ocean (the French coast); *Aplysina aerophoba* has its known distribution is in the Mediterranean Sea, and part of the South Atlantic Ocean (the Azores, the Canary Islands, and Cape Verde); *Cliona viridis* has its known distribution in the Mediterranean Sea, part of the South Atlantic Ocean (the Spanish and Portuguese coast, the Azores, the Canary Islands, and Cape Verde); *Hemimycale columella* has its known distribution in the Mediterranean Sea, part of the South Atlantic Ocean (the Spanish and Portuguese coast, the Azores, the Canary Islands, and Cape Verde), part of the North Sea (the Swedish west coast), and the Celtic Sea, including the English Channel; and finally, *Crambe crambe* has its known distribution in the Mediterranean Sea, and part of the South Atlantic Ocean (the Spanish and Portuguese coast). Source for the species range distribution: World Porifera Database.

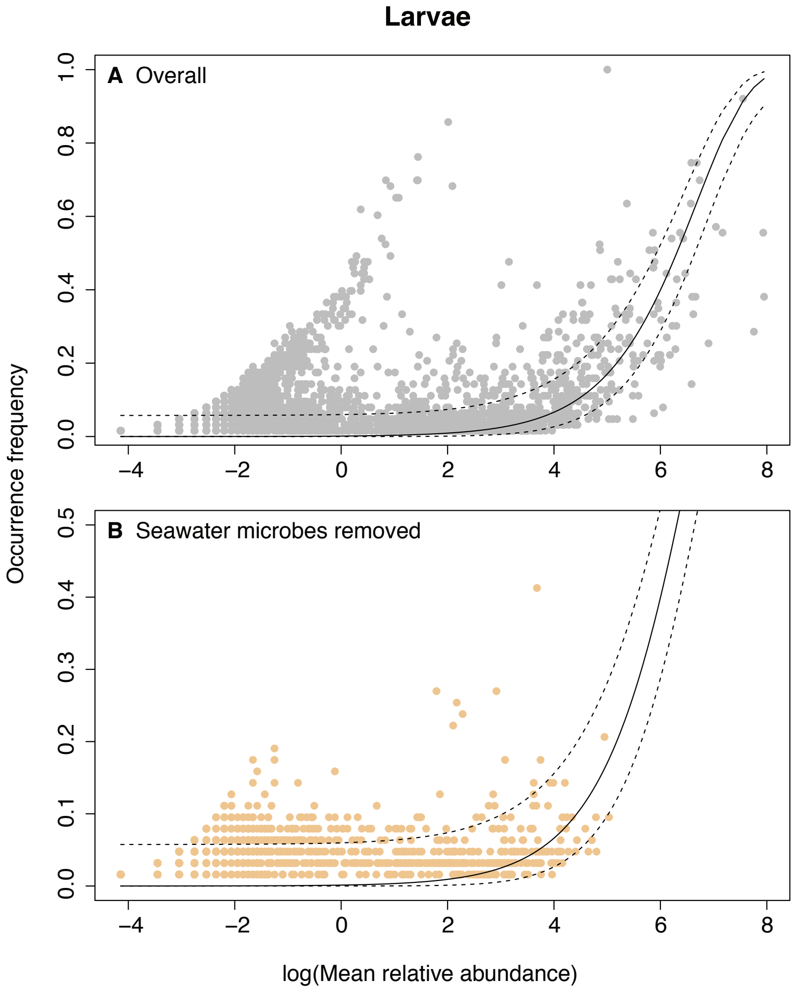


Figure S2: A “peak” consisting of taxa above the neutral prediction across individual larvae. In panel A, gray dots correspond to the overall larval microbiota. In the bottom panel B, yellow dots correspond to microbes not detected in seawater. When microbes detected in seawater are removed, much of the “peak” largely disapears.

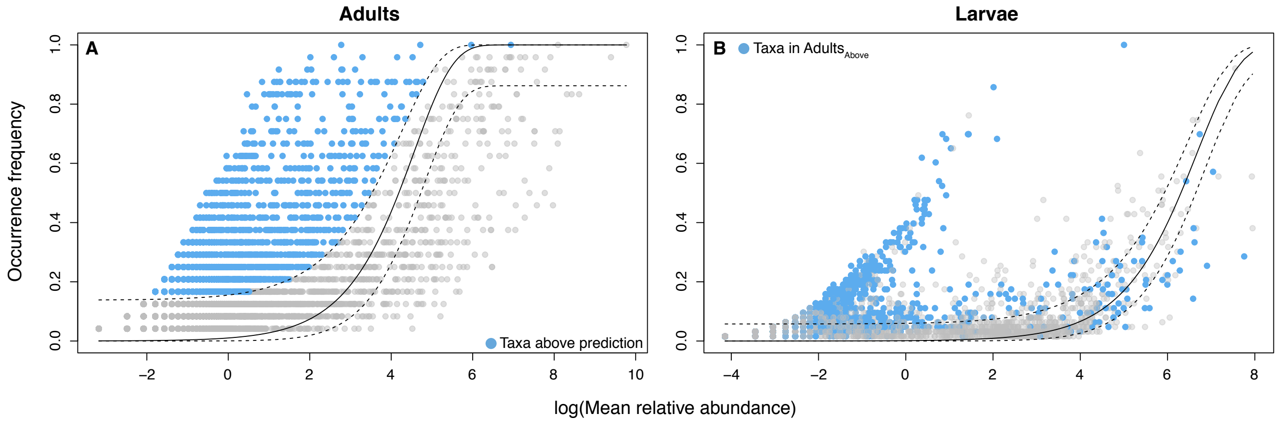


Figure S3: Most of the microbes forming the "peak" in the larval (B) are also present above the neutral prediction in adults (A). In panel A, microbes that fall above the neutral prediction in adults are colored blue. In panel B, the same microbes that fall above the neutral prediction in adults that are present in larvae are colored blue.

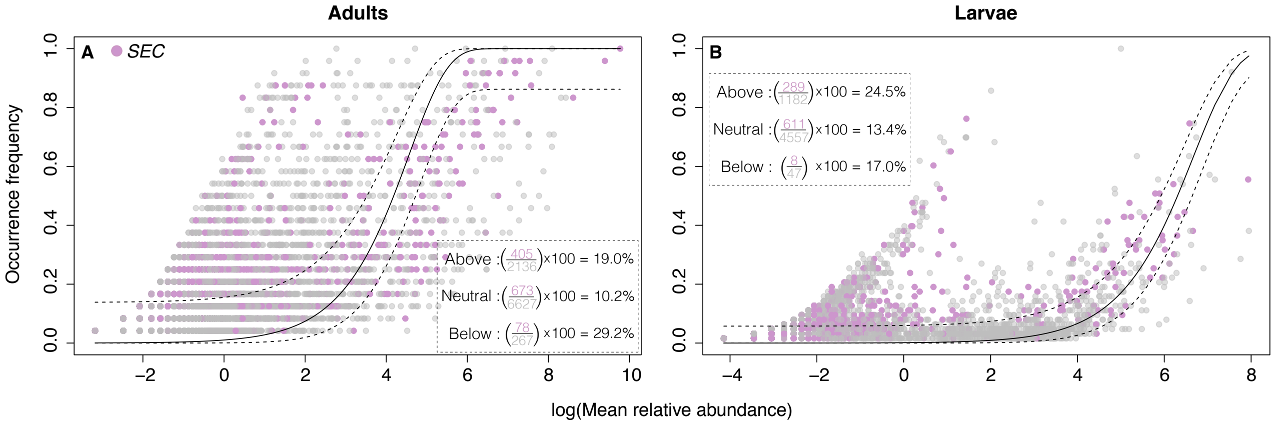


Figure S4: The distribution of *sponge-enriched clusters* (pink dots) above, within and below the neutral prediction for the (A) adults and (B) larvae. Gray dots correspond to ASVs that does not assign to *sponge-enriched clusters*. For each partition (i.e., above, within, and below), the percentage of *sponge-enriched clusters* has been calculated.

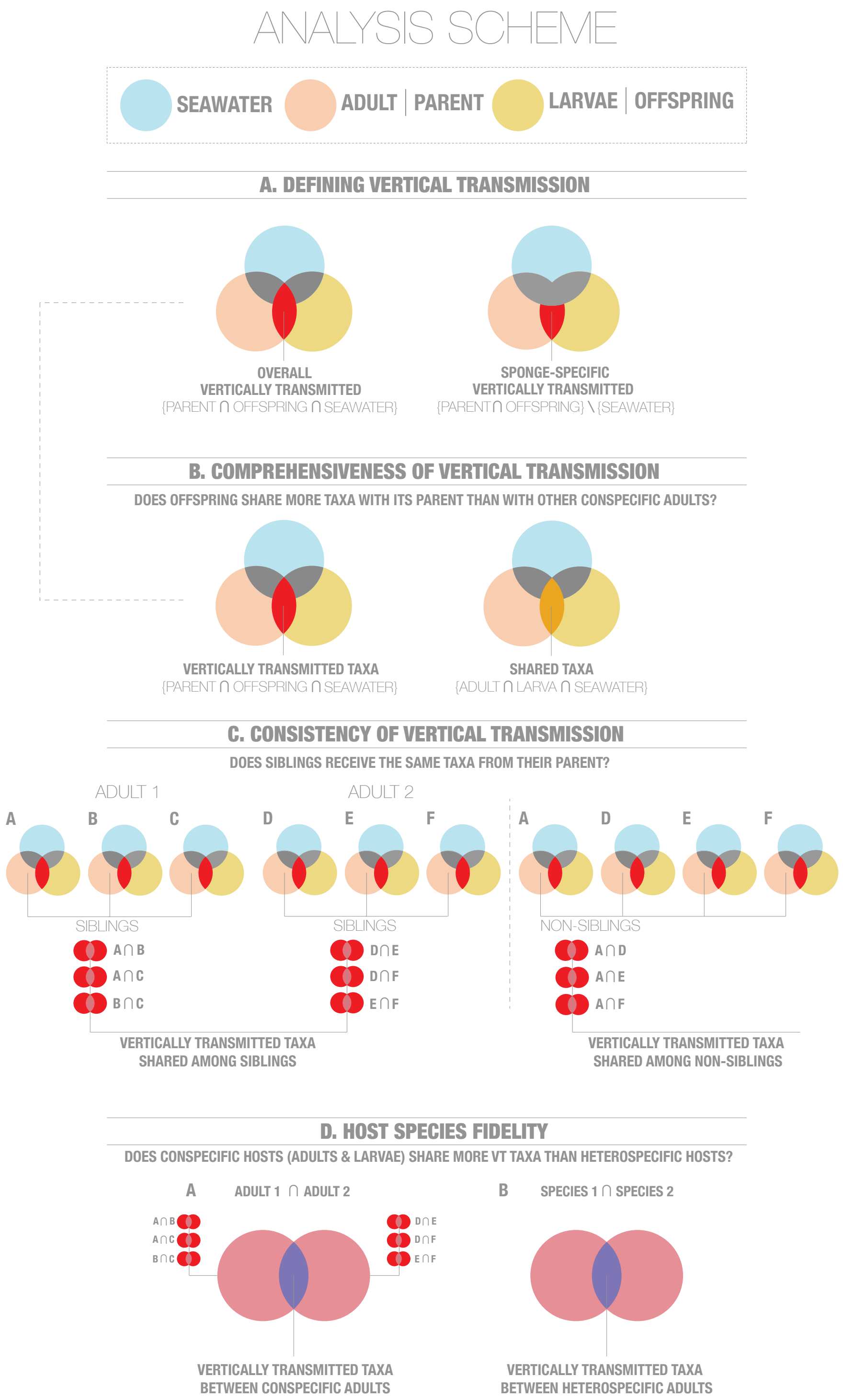
ANALYSIS SCHEME
SEAWATER
ADULT | PARENT
LARVAE | OFFSPRING
A. DEFINING VERTICAL TRANSMISSION
OVERALL
VERTICALLY TRANSMITTED
{PARENT ∩ OFFSPRING ∩ SEAWATER}
SPONGE-SPECIFIC
VERTICALLY TRANSMITTED
{PARENT ∩ OFFSPRING} \ {SEAWATER}
B. COMPREHENSIVENESS OF VERTICAL TRANSMISSION
DOES OFFSPRING SHARE MORE TAXA WITH ITS PARENT THAN WITH OTHER CONSPECIFIC ADULTS?
VERTICALLY TRANSMITTED TAXA
{PARENT ∩ OFFSPRING ∩ SEAWATER}
SHARED TAXA
{ADULT ∩ LARVA ∩ SEAWATER}
C. CONSISTENCY OF VERTICAL TRANSMISSION
DOES SIBLINGS RECEIVE THE SAME TAXA FROM THEIR PARENT?
ADULT 1
ADULT 2
A
B
C
D
E
F
A
D
E
F
SIBLINGS
SIBLINGS
NON-SIBLINGS
A ∩ B
A ∩ C
B ∩ C
D ∩ E
D ∩ F
E ∩ F
A ∩ D
A ∩ E
A ∩ F
VERTICALLY TRANSMITTED TAXA
SHARED AMONG SIBLINGS
VERTICALLY TRANSMITTED TAXA
SHARED AMONG NON-SIBLINGS
D. HOST SPECIES FIDELITY
DOES CONSPECIFIC HOSTS (ADULTS & LARVAE) SHARE MORE VT TAXA THAN HETEROSPECIFIC HOSTS?
A
ADULT 1 ∩ ADULT 2
B
SPECIES 1 ∩ SPECIES 2
A ∩ B
A ∩ C
B ∩ C
D ∩ E
D ∩ F
E ∩ F
VERTICALLY TRANSMITTED TAXA
BETWEEN CONSPECIFIC ADULTS
VERTICALLY TRANSMITTED TAXA
BETWEEN HETEROSPECIFIC ADULTS

Figure S5

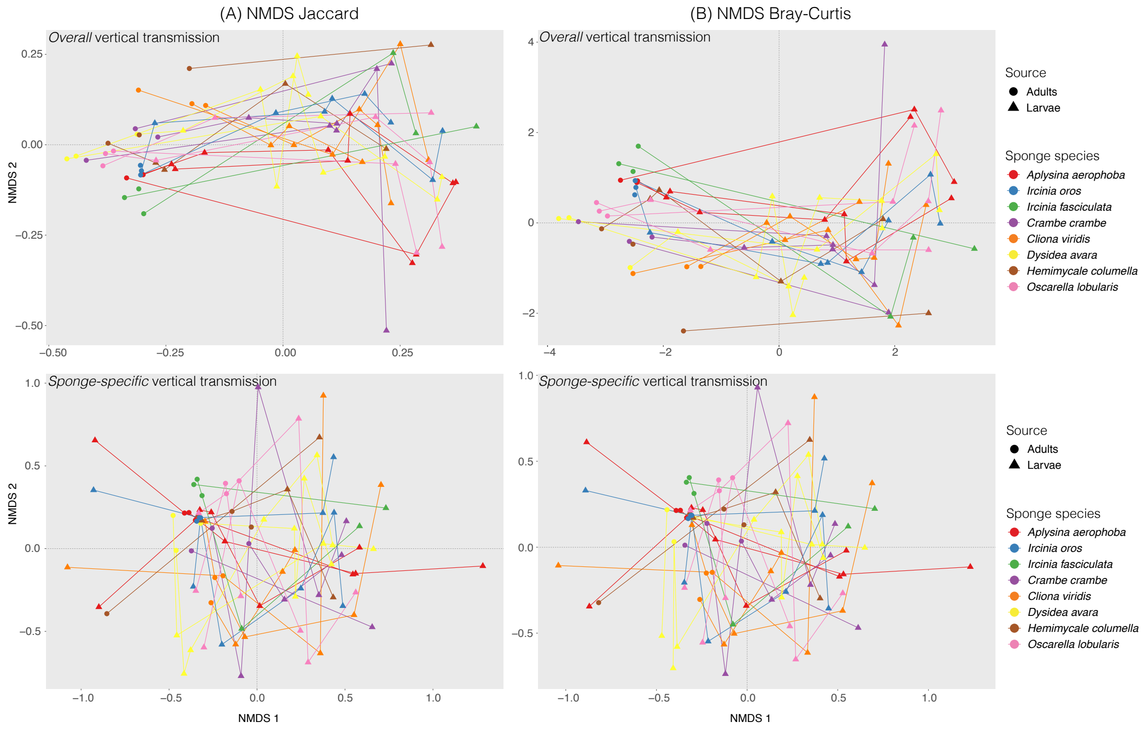


Figure S6: Non metric multidimensional scaling (NMDS) for Jaccard (A) and Bray-Curtis (B) distances calculated for the *overall* and *sponge-specific* definition of vertical transmission, respectively.

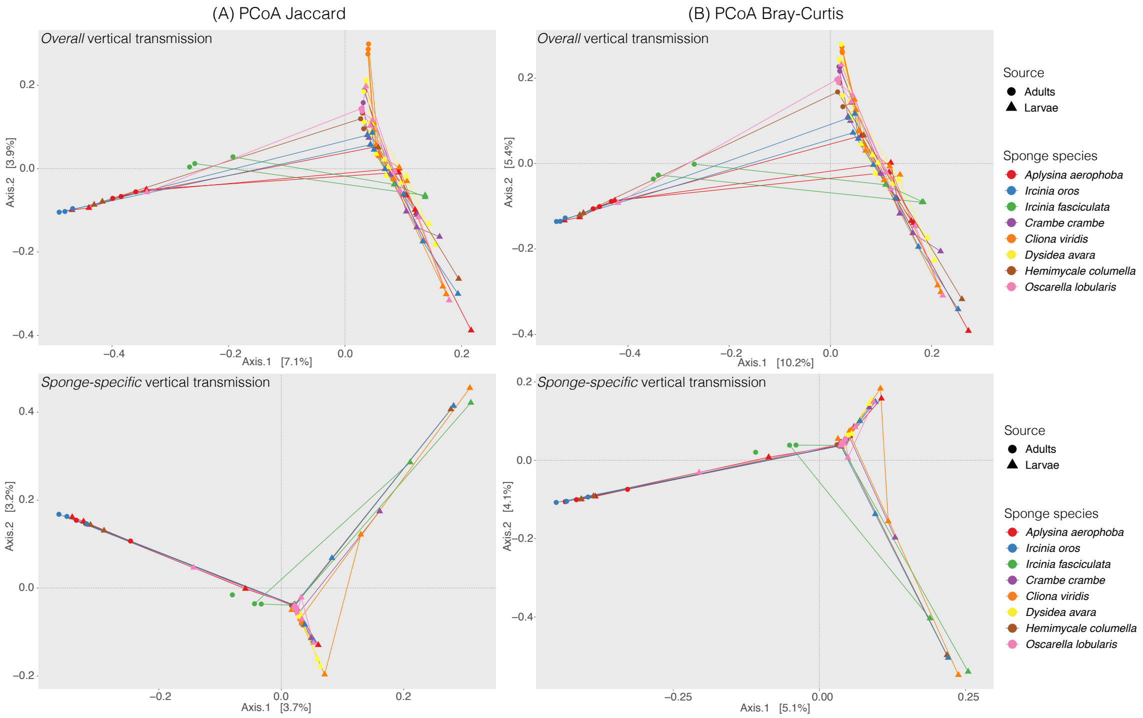


Figure S7: Principle Coordinates Analysis (PCoA) for Jaccard (A) and Bray-Curtis (B) distances calculated for the *overall* and *sponge-specific* definition of vertical transmission, respectively.

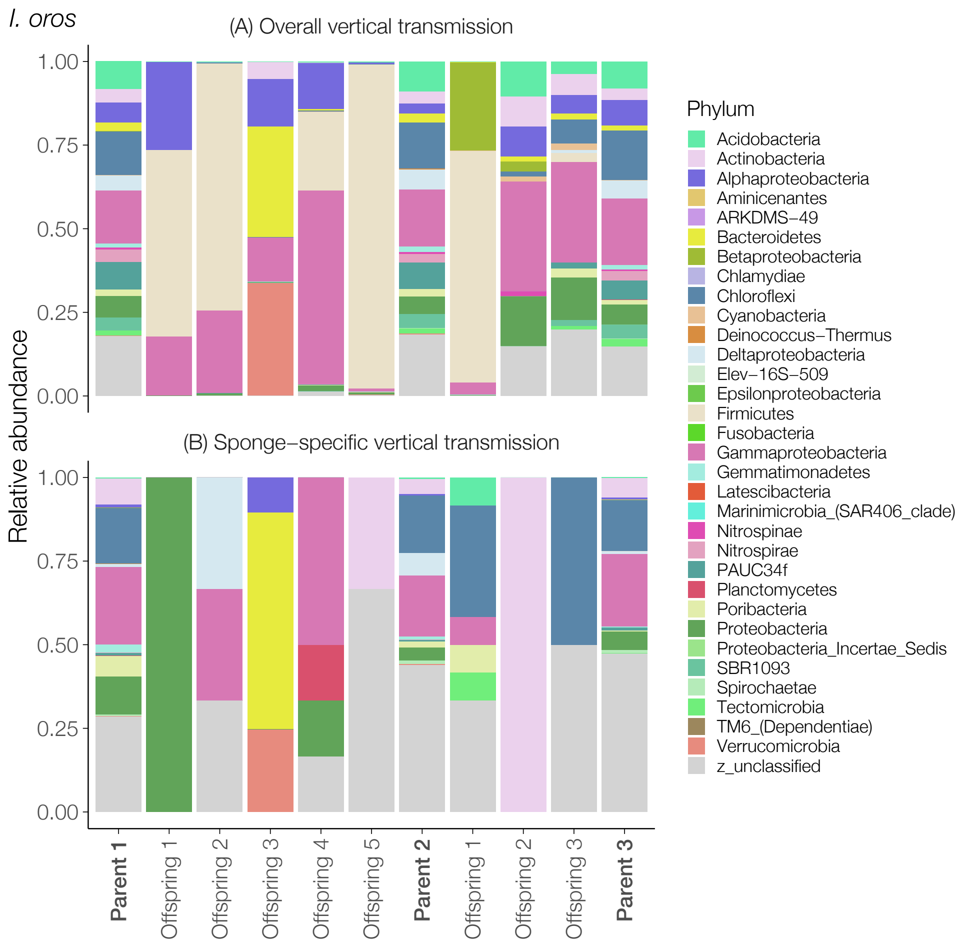


a. See figure legend below

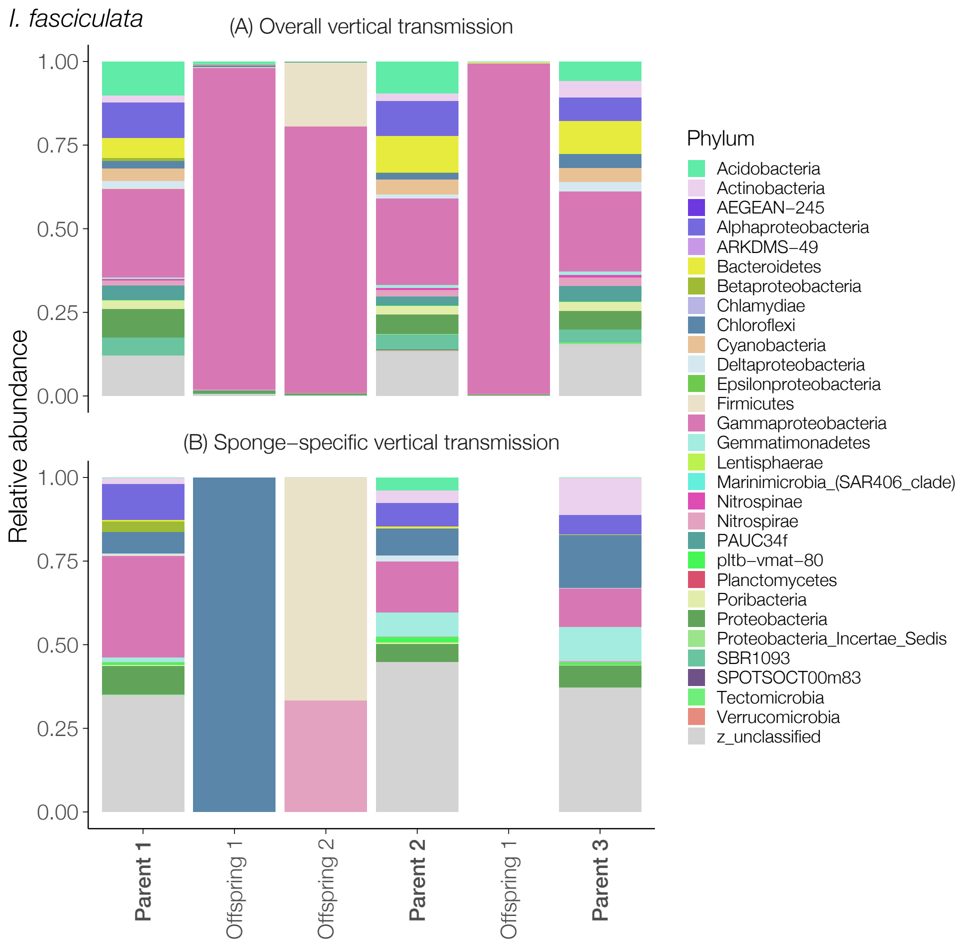


b. See figure legend below

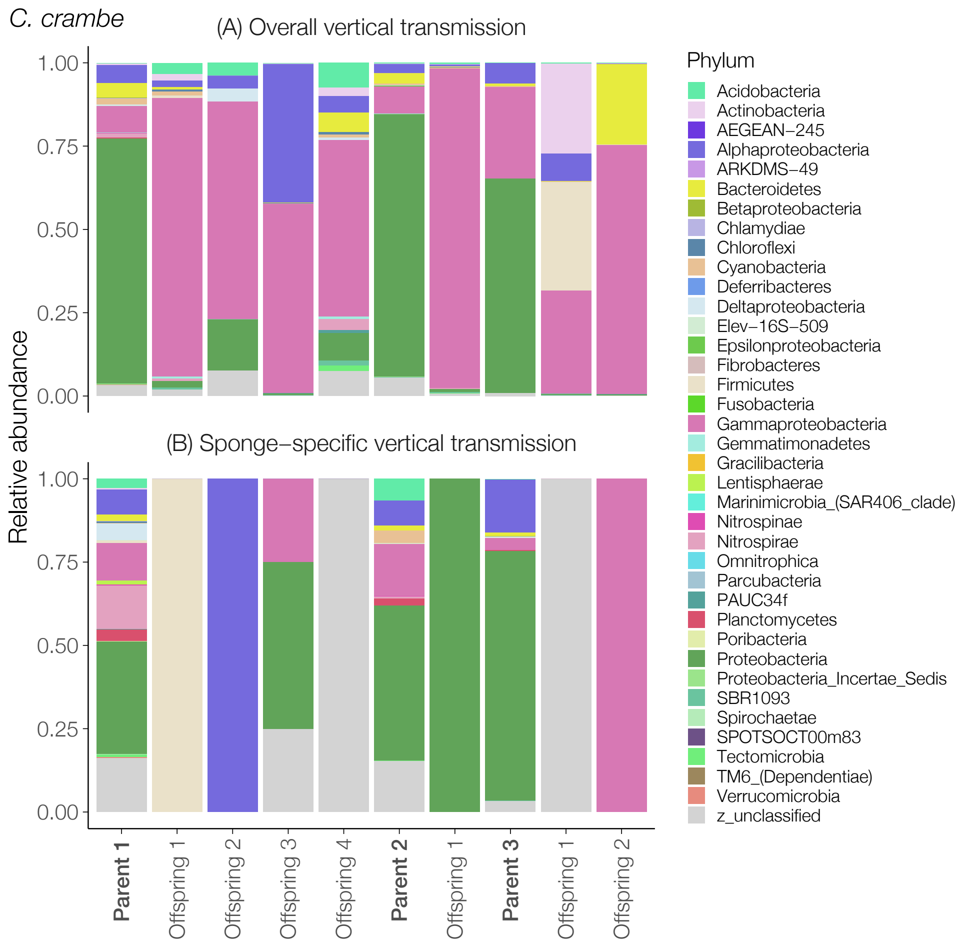


c. See figure legend below

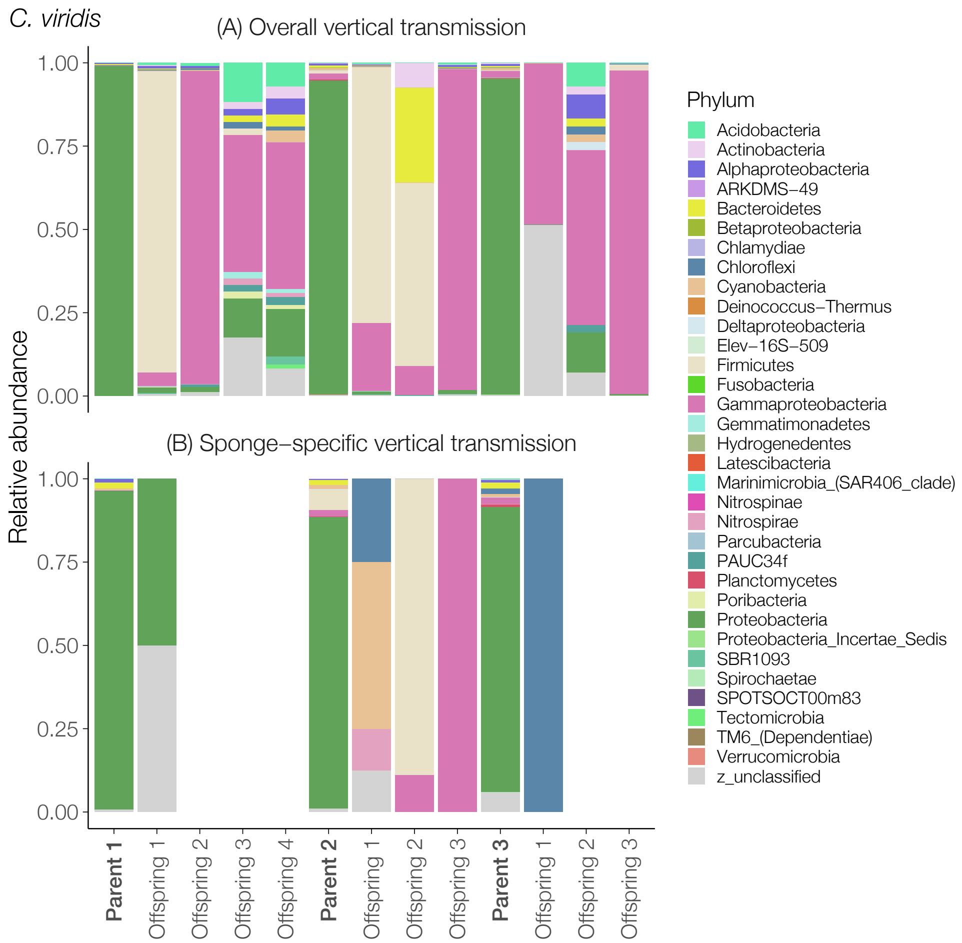


d. See figure legend below

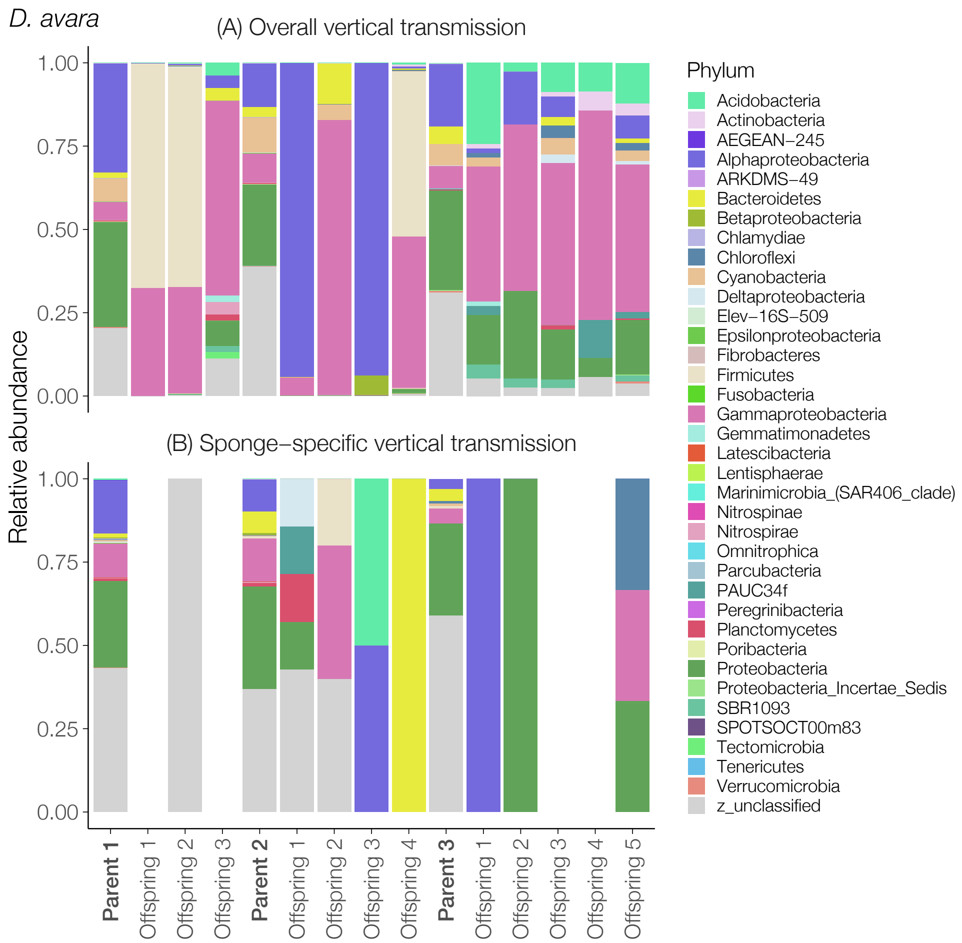


e. See figure legend below

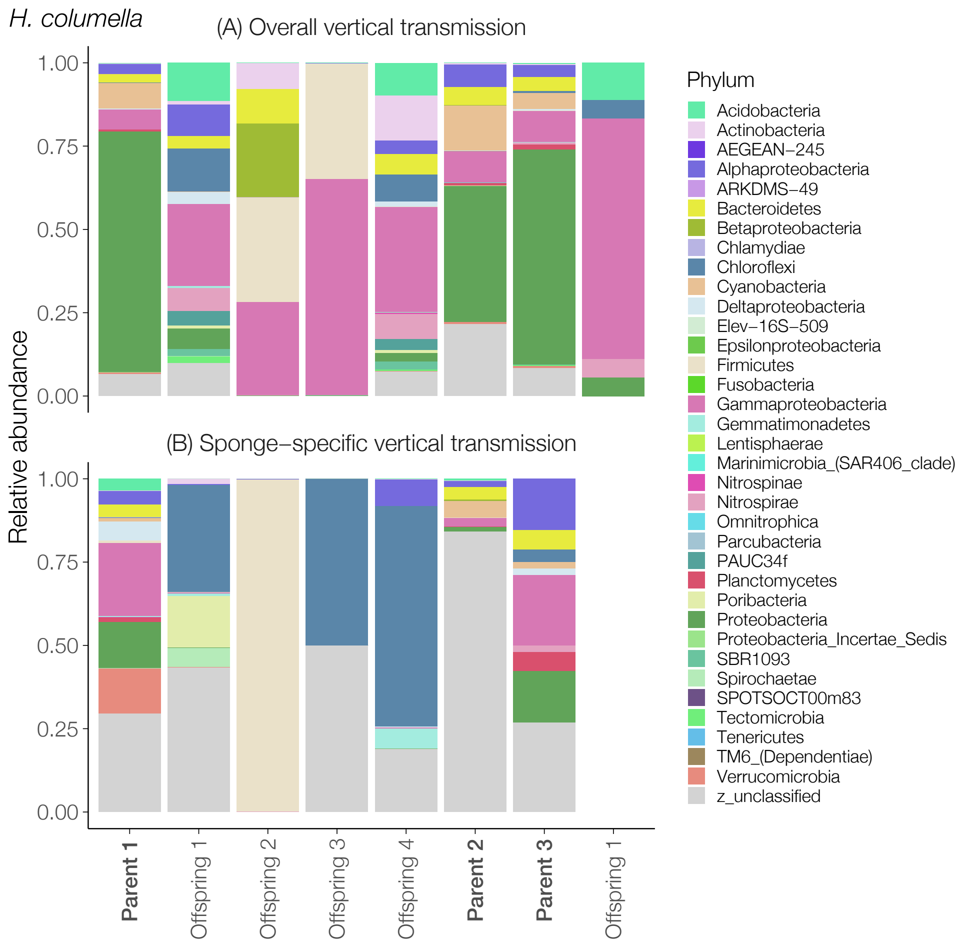


f. See figure legend below

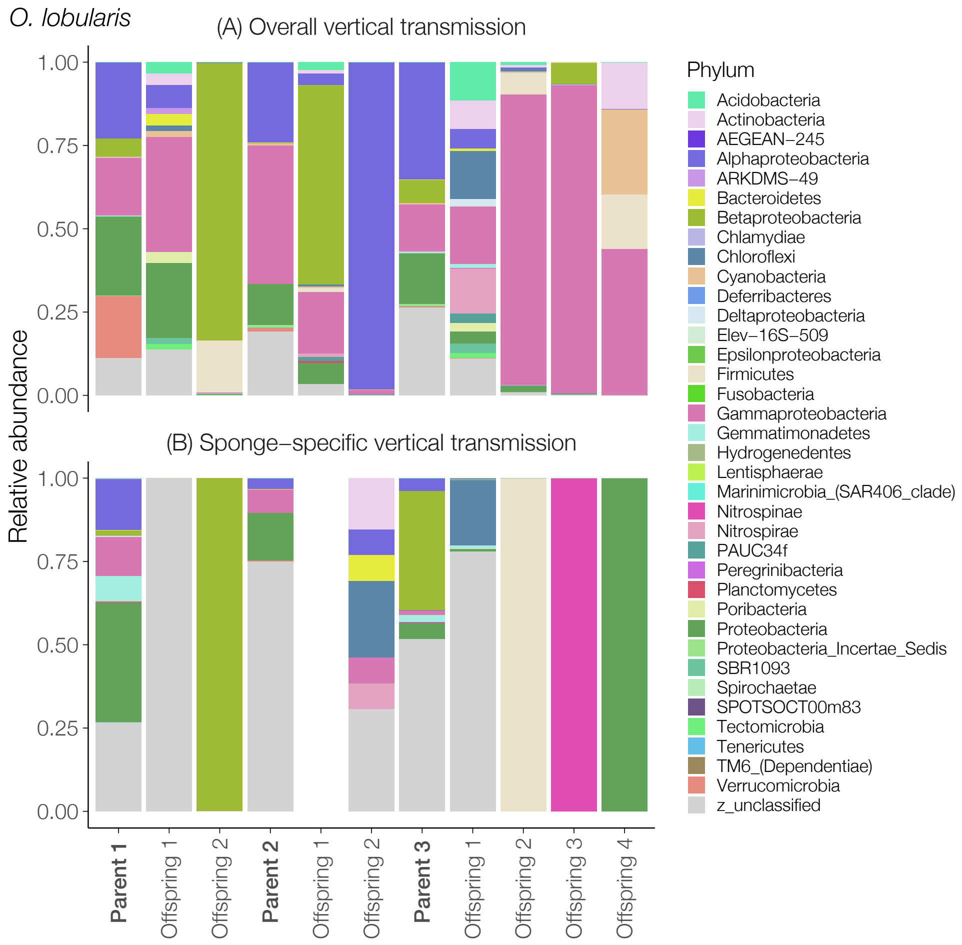


g. See figure legend below

Figure S8: The relative contribution of vertically transmitted ASVs classifying to different microbial phyla (classes for Proteobacteria) in parents and the offspring of the remaining seven sponge species. The top panel (A) shows the relative contribution of phyla for the *overall* definition of vertical transmission, and the bottom panel (B) shows the relative contribution of phyla for *sponge-specific* vertical transmission. Parents (in bold) and offspring are shown on the x-axis. Note that when microbes detected in seawater are removed, this sometimes leaves no vertical transmitted ASVs for the *sponge-specific* vertical transmission. Colors represent different microbial phyla (classes for Proteobacteria). Panels a-g corresponds to host species: *I. oros; I. fasciculata; C. crambe; C. viridis; D. avara; H. columella*; and *O. lobularis*.

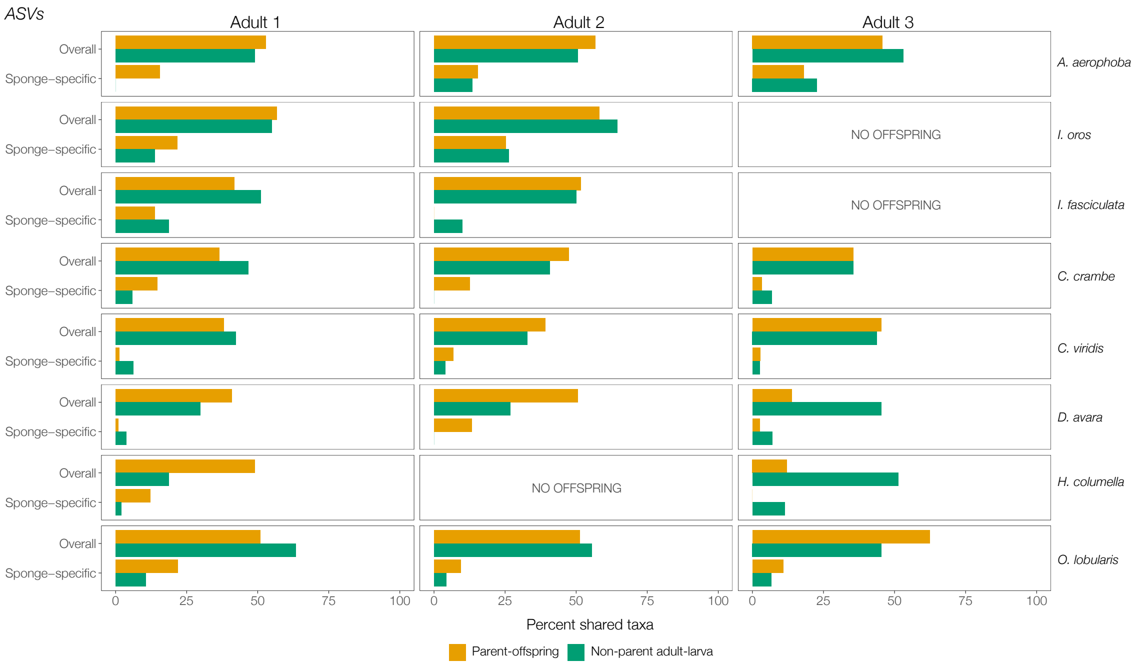


Figure S9: The average percent shared ASVs in the (A) *overall* and (B) *sponge-specific* definition of vertical transmission across adults and sponge species. Barplots show the average percent shared ASVs between sponge larvae and either (i) their known parents (yellow bars), or (ii) non-parental conspecific adults (green bars). Each row corresponds to a sponge species, and each column one of the three adults for that focal species.

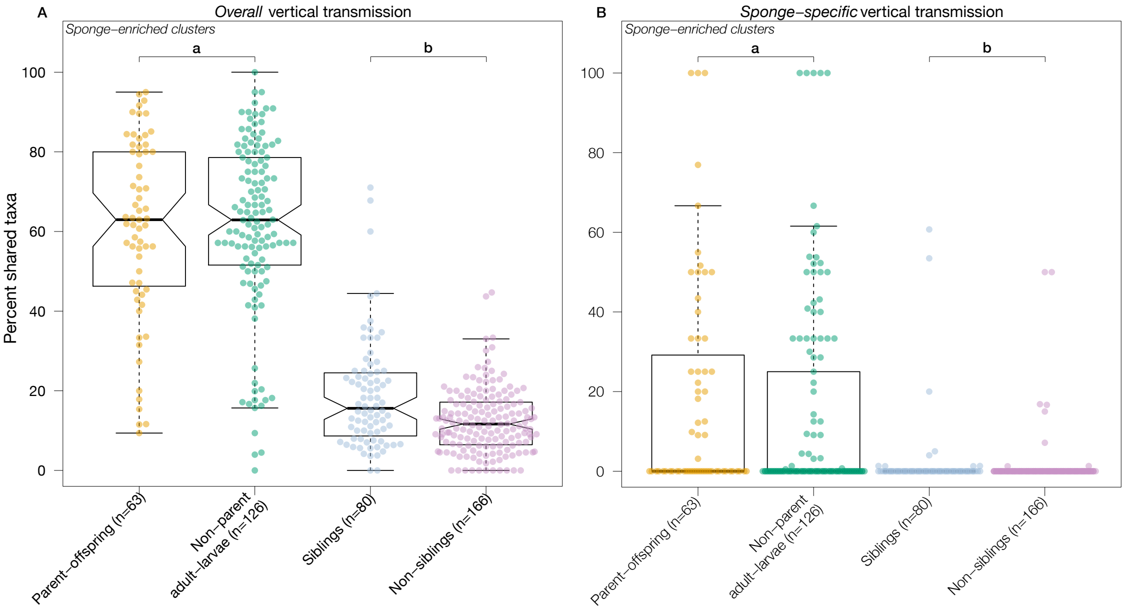


Figure S10: Percent shared *sponge-enriched clusters* in the (A) *overall* and (B) *sponge-specific* definition of vertical transmission. Boxplots (a) show the percent shared *sponge-enriched clusters* between sponge larvae and either (i) their known parents (yellow dots), or (ii) non-parental conspecific adults (green dots). In boxplots (a), each dot represents one parent-offspring pair, or one non-parent adult-larva pair across all sponge species (see Figure S11). For *overall* vertical transmission (A), parents and offspring shared, on average, 60.7% of the ASVs, whereas non-parental conspecific adults and larvae shared, on average, 61.3% of the *sponge-enriched clusters* (P>0.1). For *sponge-specific* vertical transmission (B), parents and offspring shared, on average, 18.6% of the *sponge-enriched clusters*, whereas non-parental conspecific adults and larvae shared, on average, 14.12% of the *sponge-enriched clusters* (P>0.1). Boxplots (b) show the percent shared vertically transmitted *sponge-enriched clusters* between (i) siblings (blue dots), and (ii) non-siblings (purple dots). In boxplots (b), each dot represents one sibling pair, or one pair of non-siblings (see Figure S12). For *overall* vertical transmission (A), siblings shared, on average, 18.8% of their vertically transmitted ASVs, while non-siblings only shared 12.4% % (P<0.001). For *sponge-specific* vertical transmission (B), siblings shared, on average, only 1.85% of their vertically transmitted ASVs, whereas non-siblings shared 0.6% (P=0.024). While these are significantly different, the effect size (i.e., the difference in location, Δ), is effectively zero.

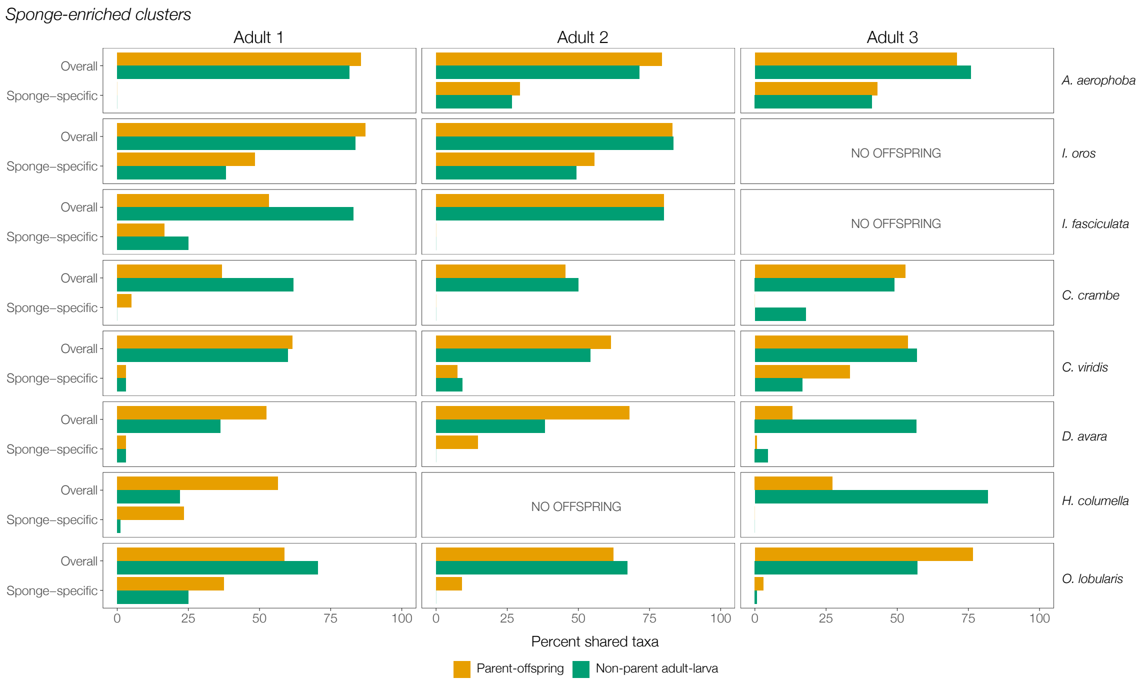


Figure S11: The average percent shared *sponge-enriched clusters* in the (A) *overall* and (B) *sponge-specific* definition of vertical transmission across adults and sponge species. Barplots show the average percent shared *sponge-enriched clusters* between sponge larvae and either (i) their known parents (yellow bars), or (ii) non-parental conspecific adults (green bars). Each row corresponds to a sponge species, and each column one of the three adults for that focal species.

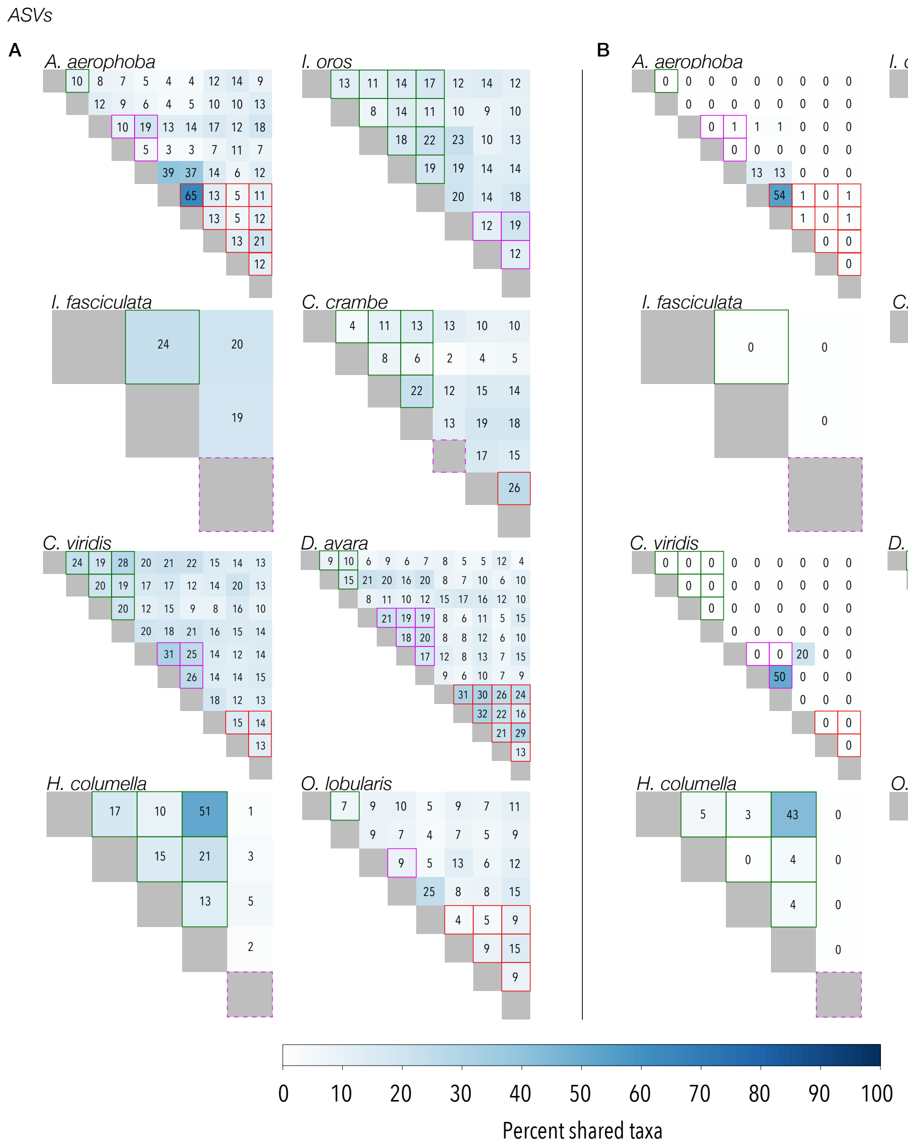


Figure S12: Percent shared ASVs of (A) *overall* and (B) *sponge-specific* vertical transmission between siblings and non-siblings. Each cell represents a larva, and sets of siblings from the same parent are indicated by cells bordered by the same color (green, purple, or red). In cases where parents only had one offspring, the diagonal is bordered by a dashed line. Cells with no boarders correspond to the percent of vertically transmitted ASVs that are shared between non-siblings, i.e., conspecific larvae that did not share the same parent. Solid gray cells (i.e., the diagonal) represent the comparison with self. The white-blue continuous color legend corresponds to 0% (no ASVs shared) in white and 100% (all ASVs shared) in dark blue).

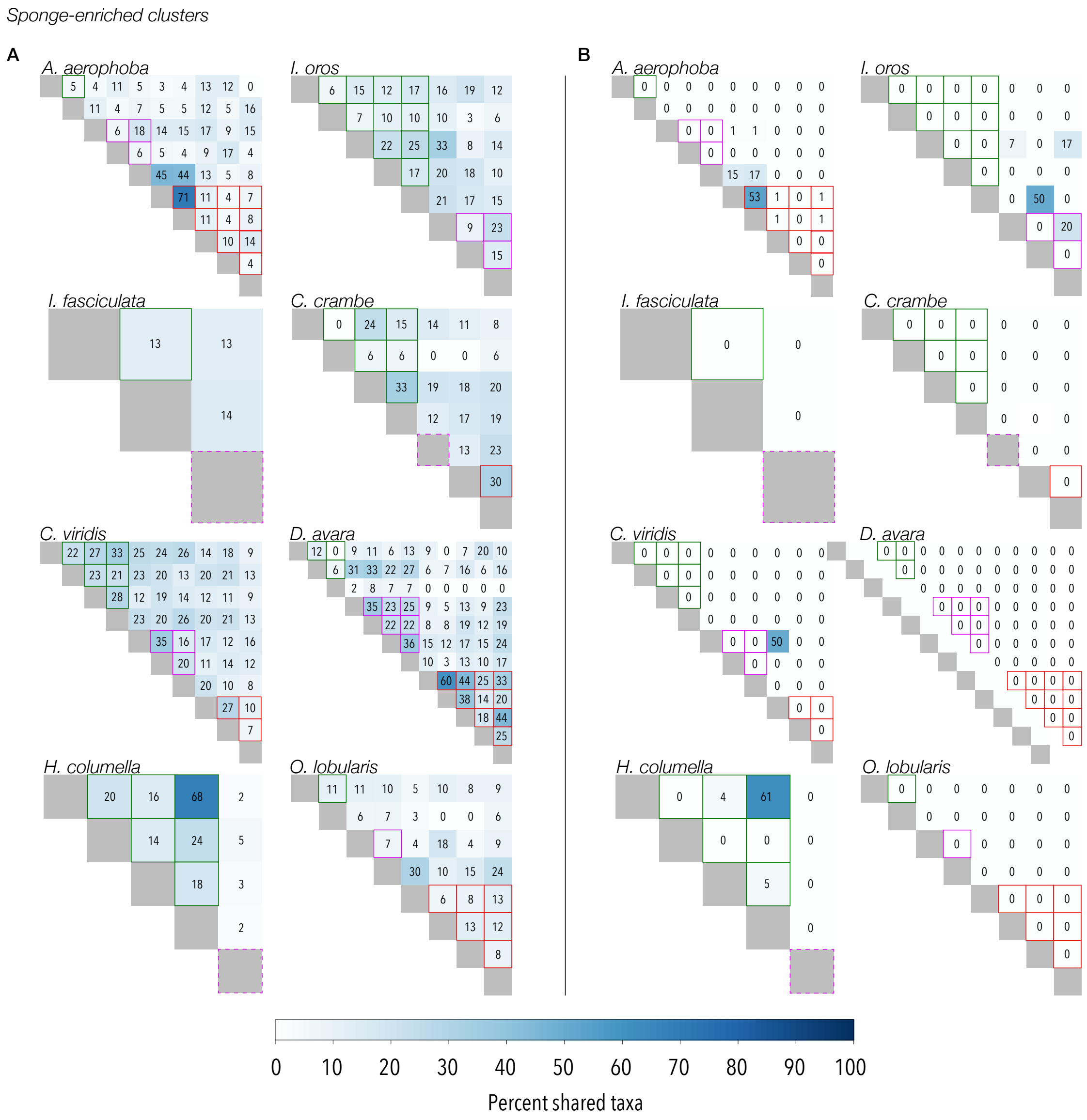


Figure S13: Percent shared *sponge-enriched clusters* of (A) *overall* and (B) *sponge-specific* vertical transmission between siblings and non-siblings. Each cell represents a larva, and sets of siblings from the same parent are indicated by cells bordered by the same color (green, purple, or red). In cases where parents only had one offspring, the diagonal is bordered by a dashed line. Cells with no boarders correspond to the percent of vertically transmitted ASVs that are shared between non-siblings, i.e., conspecific larvae that did not share the same parent. Solid gray cells (i.e., the diagonal) represent the comparison with self. The white-blue continuous color legend corresponds to 0% (no *sponge-enriched clusters* shared) in white and 100% (all *sponge-enriched clusters* shared) in dark blue).

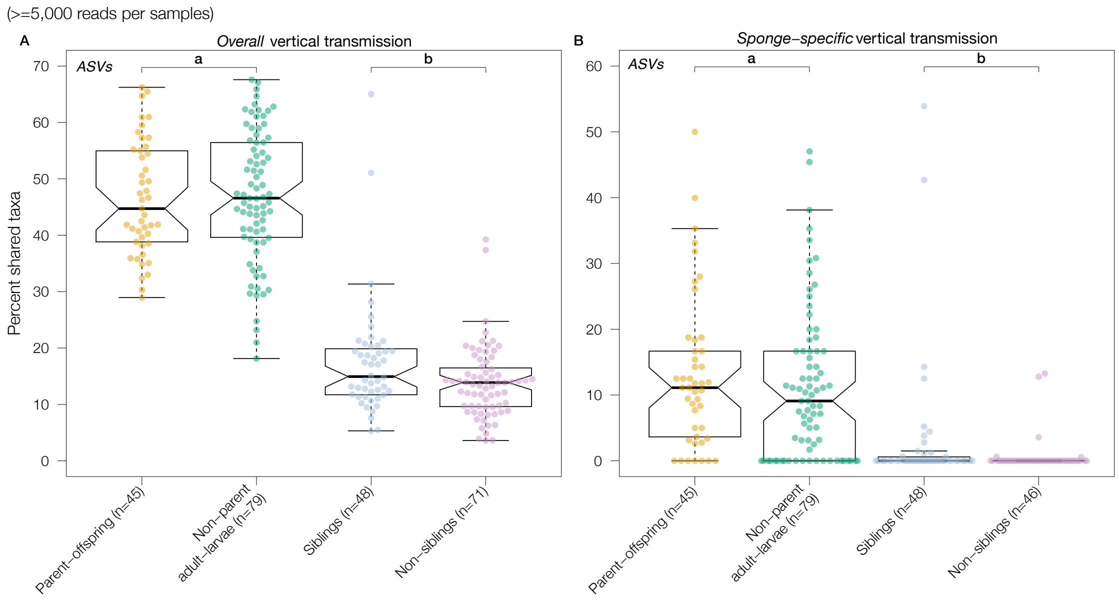


Figure S14: Percent shared ASVs in the (A) *overall* and (B) *sponge-specific* definition of vertical transmission for samples with at least 5,000 sequence reads. Boxplots (a) show the percent shared ASVs between sponge larvae and either (i) their known parents (yellow dots), or (ii) non-parental conspecific adults (green dots). In boxplots (a), each dot represents one parent-offspring pair, or one non-parent adult-larva pair across all sponge species. For *overall* vertical transmission (A), parents and offspring shared, on average, 46.5% of the ASVs, whereas non-parental conspecific adults and larvae shared, on average, 46.8% of the ASVs (Δ=-1.09, 95% CI [-5.18,3.26], Mann-Whitney U=1688.5, P>0.1). For *sponge-specific* vertical transmission (B), parents and offspring shared, on average, 13.1% of the ASVs, whereas non-parental conspecific adults and larvae shared, on average, 11.1% of the ASVs (Δ=1.79, 95% CI [-0.76,5.20], Mann-Whitney U=2006.5, P>0.1). Boxplots (b) show the percent shared VT ASVs between (i) siblings (blue dots), and (ii) non-siblings (purple dots). In boxplots (b), each dot represents one sibling pair, or one pair of non-siblings. For *overall* vertical transmission (A), siblings shared, on average, 17.5% of their VT ASVs, while non-siblings only shared 13.8% (Δ=2.66, 95% CI [0.23,4.83], Mann-Whitney U=2101.5, P=0.03). For *sponge-specific* vertical transmission (B), siblings shared, on average, only 3.0% of their VT ASVs, whereas non-siblings shared 1.3% (Δ=0.00, 95% CI [-0.00,0.00], Mann-Whitney U=1947, P=0.05). While these are significantly different, the effect size (i.e., the difference in location, Δ), is effectively zero.

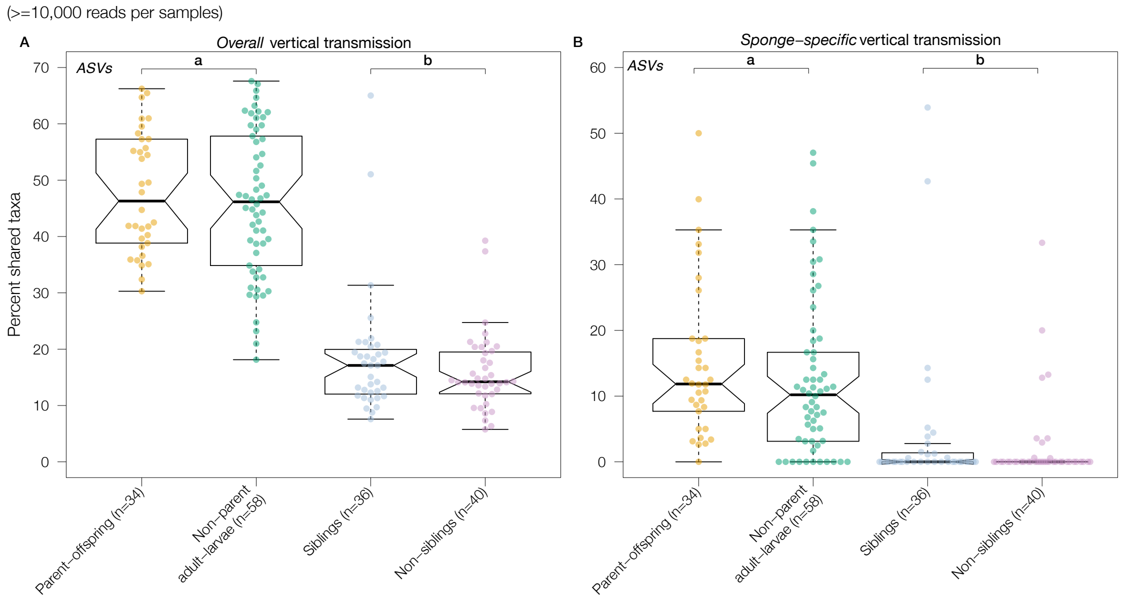


Figure S15: Percent shared ASVs in the (A) *overall* and (B) *sponge-specific* definition of vertical transmission for samples with at least 10,000 sequence reads. Boxplots (a) show the percent shared ASVs between sponge larvae and either (i) their known parents (yellow dots), or (ii) non-parental conspecific adults (green dots). In boxplots (a), each dot represents one parent-offspring pair, or one non-parent adult-larva pair across all sponge species. For *overall* vertical transmission (A), parents and offspring shared, on average, 47.5% of the ASVs, whereas non-parental conspecific adults and larvae shared, on average, 45.9% of the ASVs ($\Delta$=1.29, 95% CI [-4.03,7.21], Mann-Whitney U=1044.5, P>0.1). For *sponge-specific* vertical transmission (B), parents and offspring shared, on average, 15.0% of the ASVs, whereas non-parental conspecific adults and larvae shared, on average, 12.4% of the ASVs ($\Delta$=2.78, 95% CI [-0.90,6.84], Mann-Whitney U=1164.5, P>0.1). Boxplots (b) show the percent shared VT ASVs between (i) siblings (blue dots), and (ii) non-siblings (purple dots). In boxplots (b), each dot represents one sibling pair, or one pair of non-siblings. For *overall* vertical transmission (A), siblings shared, on average, 18.4% of their VT ASVs, while non-siblings only shared 15.7% ($\Delta$=1.07, 95% CI [-1.52,4.01], Mann-Whitney U=789.5, P>0.1). For *sponge-specific* vertical transmission (B), siblings shared, on average, only 4.0% of their VT ASVs, whereas non-siblings shared 2.3% ($\Delta$=0.00, 95% CI [-0.00,0.00], Mann-Whitney U=815.5, P>0.1). Likely due to the reduction in sample sizes, no significant differences were found in this analysis.

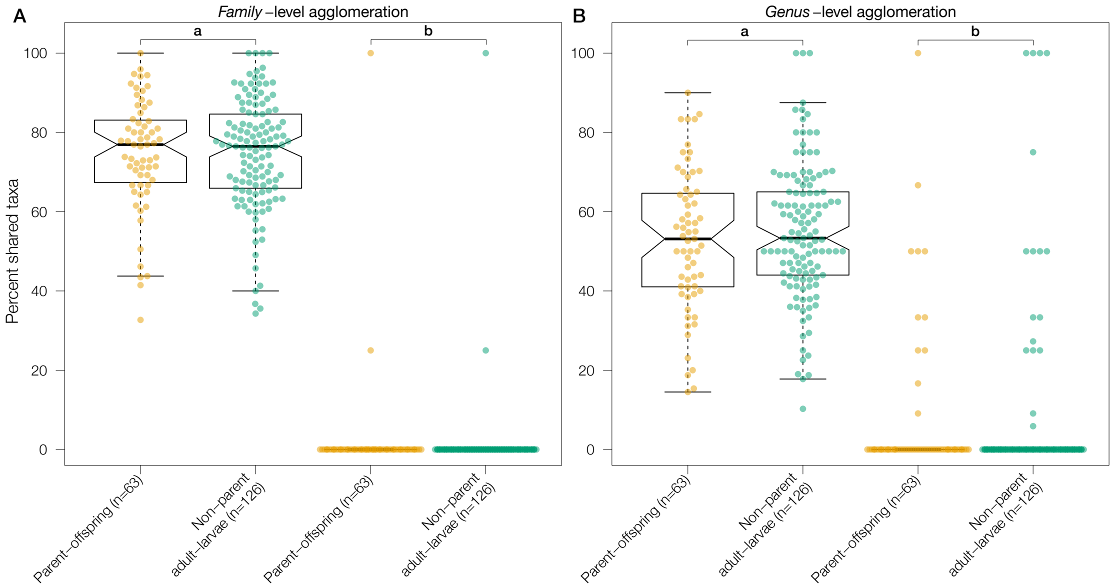


Figure S16: Percent shared taxa at the family (A) and genus (B) level in the (a) *overall* and (b) *sponge-specific* definition of vertical transmission. The boxplots show the percent shared taxa between sponge larvae and either (i) their known parents (yellow dots), or (ii) non-parental conspecific adults (green dots). Each dot represents one parent-offspring pair, or one non-parent adult-larva pair across all sponge species. At the family level (A), parents and offspring shared, on average, 74.5% of the *overall* taxa, whereas non-parental conspecific adults and larvae shared, on average, 74.7% of the *overall* taxa (Δ=0.16, 95% CI [-4.01,4.12], Mann-Whitney U=4004.5, P>0.1). Removing taxa detected in seawater reduced the sharing even further. Parents and offspring shared, on average, 2.0% of the *sponge-specific* taxa, whereas non-parental conspecific adults and larvae shared, on average, 1.0% of the *overall* taxa (Δ=0.00, 95% CI [-0.00,0.00], Mann-Whitney U=4032, P>0.1). At the genus level (B), parents and offspring shared, on average, 52.8% of the *overall* taxa, whereas non-parental conspecific adults and larvae shared, on average, 55.0% of the *overall* taxa (Δ=-1.91, 95% CI [-7.14,3.48], Mann-Whitney U=3734.5, P>0.1). Also here, removing taxa detected in seawater reduced the sharing even further. Parents and offspring shared, on average, 7.3% of the *sponge-specific* taxa, whereas non-parental conspecific adults and larvae shared, on average, 6.8% of the *overall* taxa (Δ=0.00, 95% CI [-0.00,0.00], Mann-Whitney U=4114.5, P>0.1).

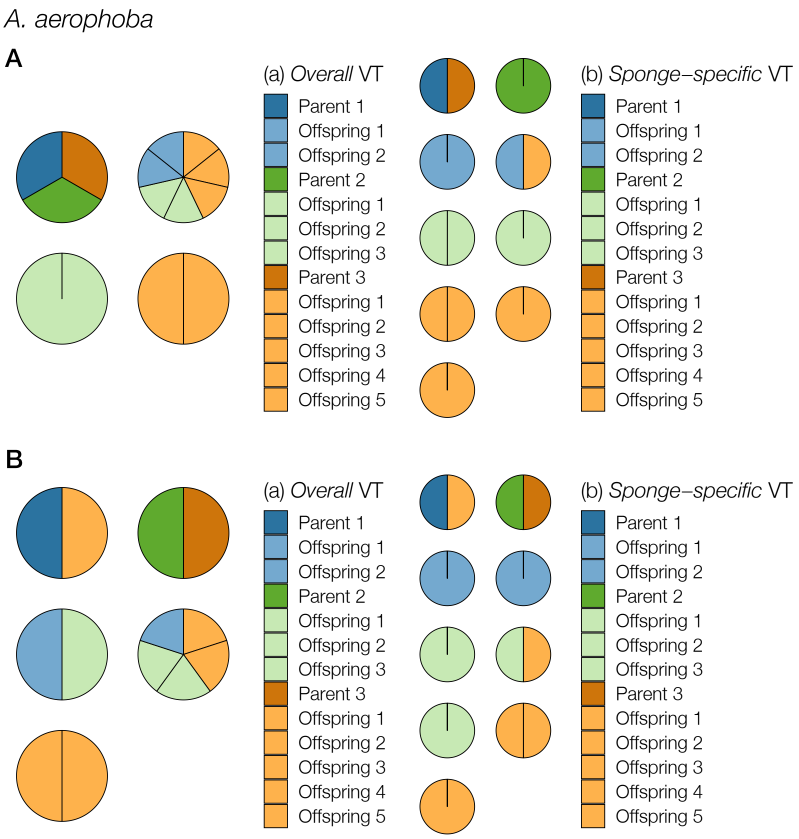


i. See figure legend below

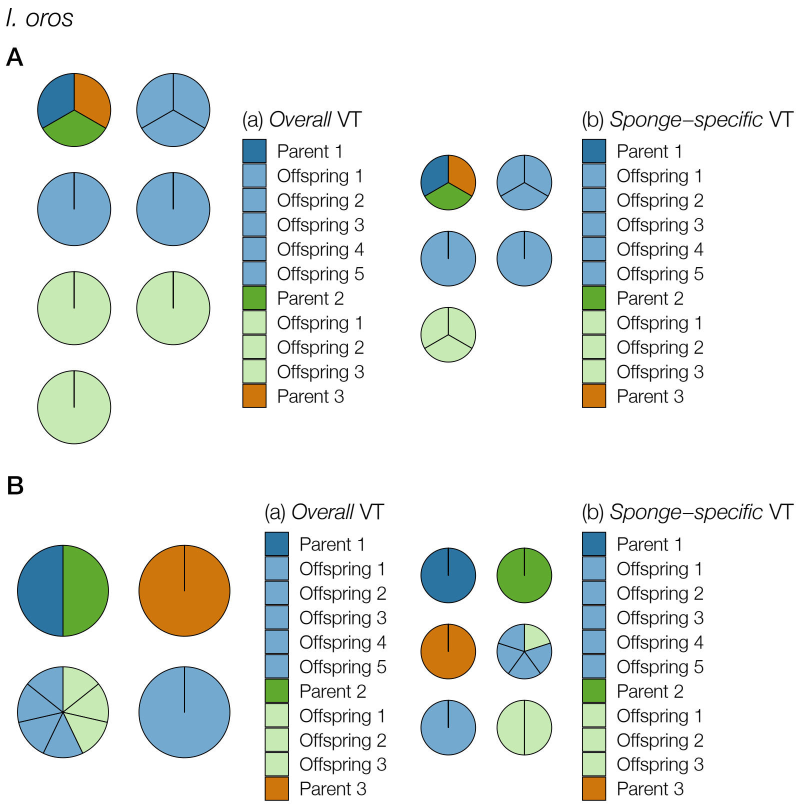


ii. See figure legend below

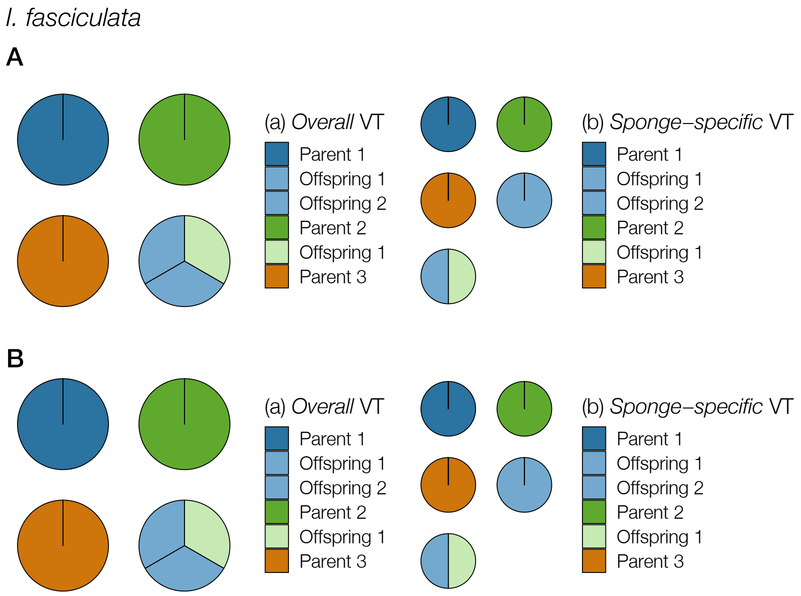


iii. See figure legend below

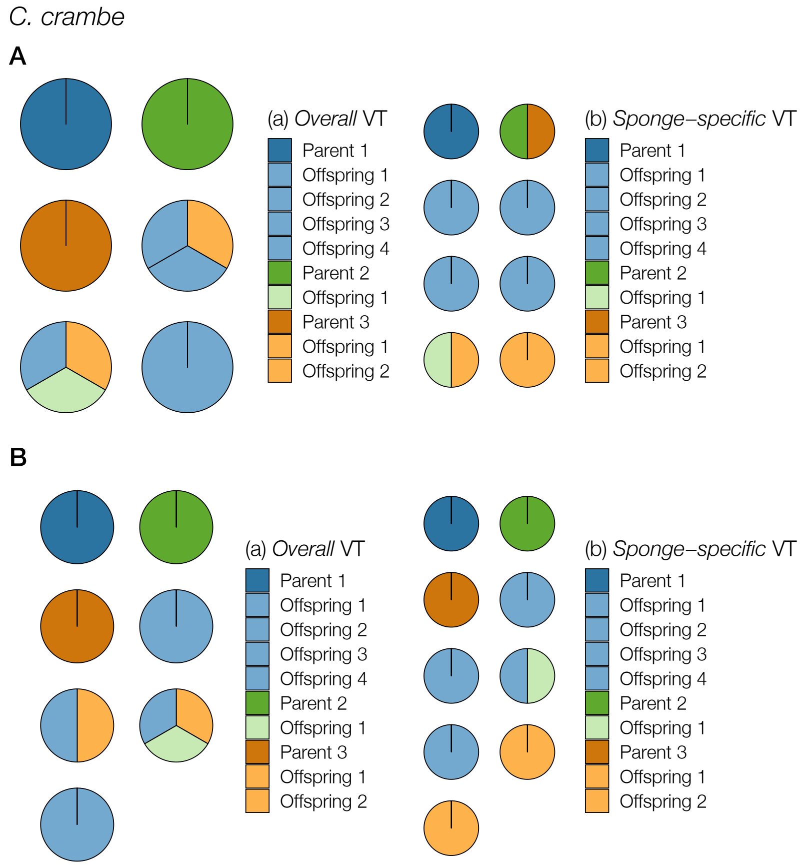


iv. See figure legend below

*C. viridis*

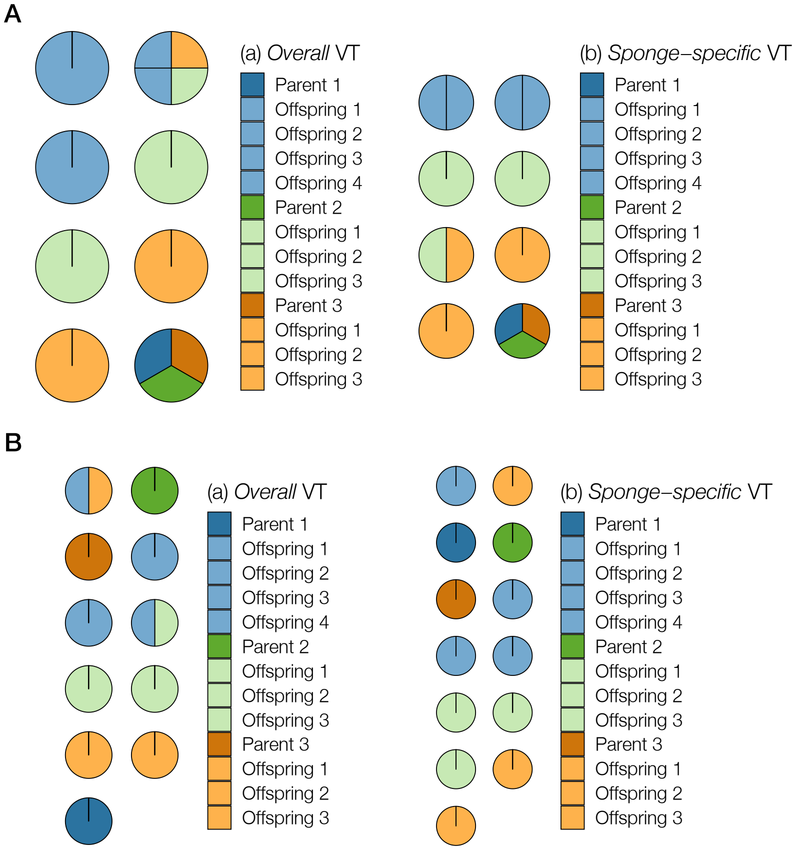


v. See figure legend below

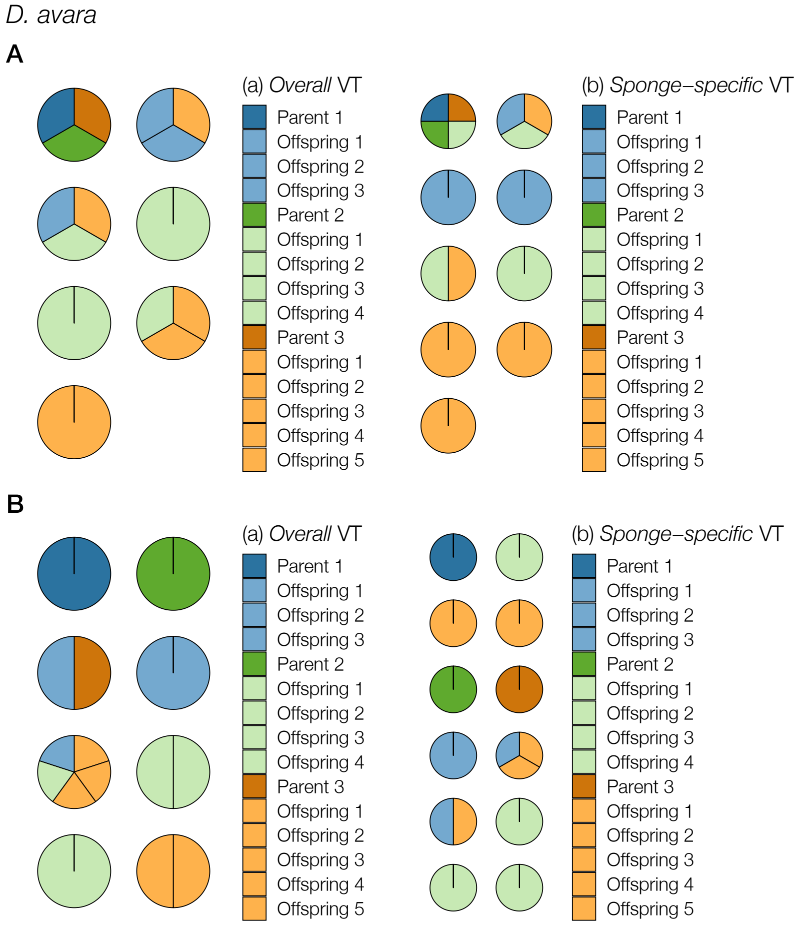


vi. See figure legend below

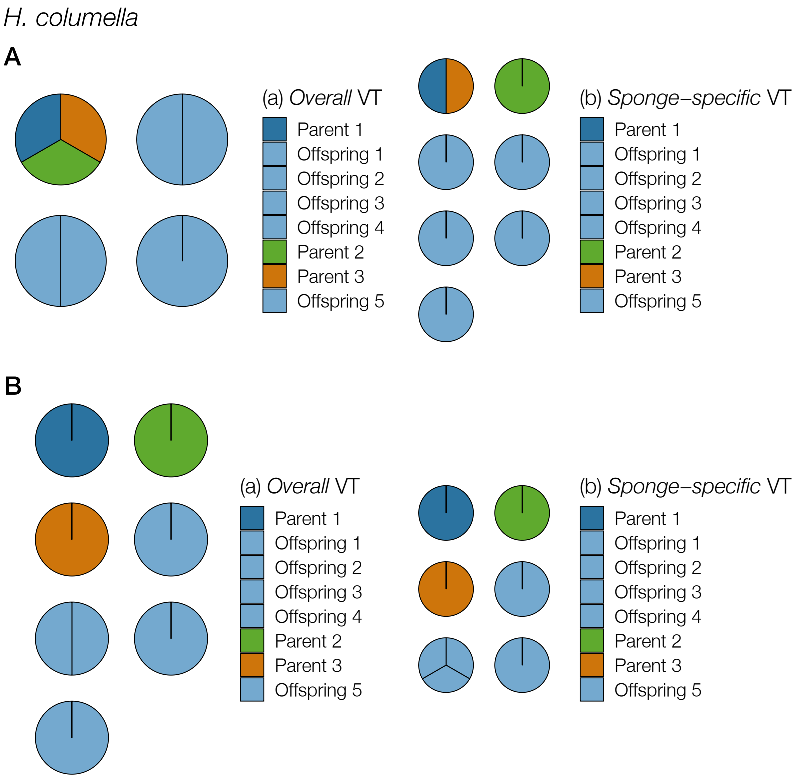


vii. See figure legend below

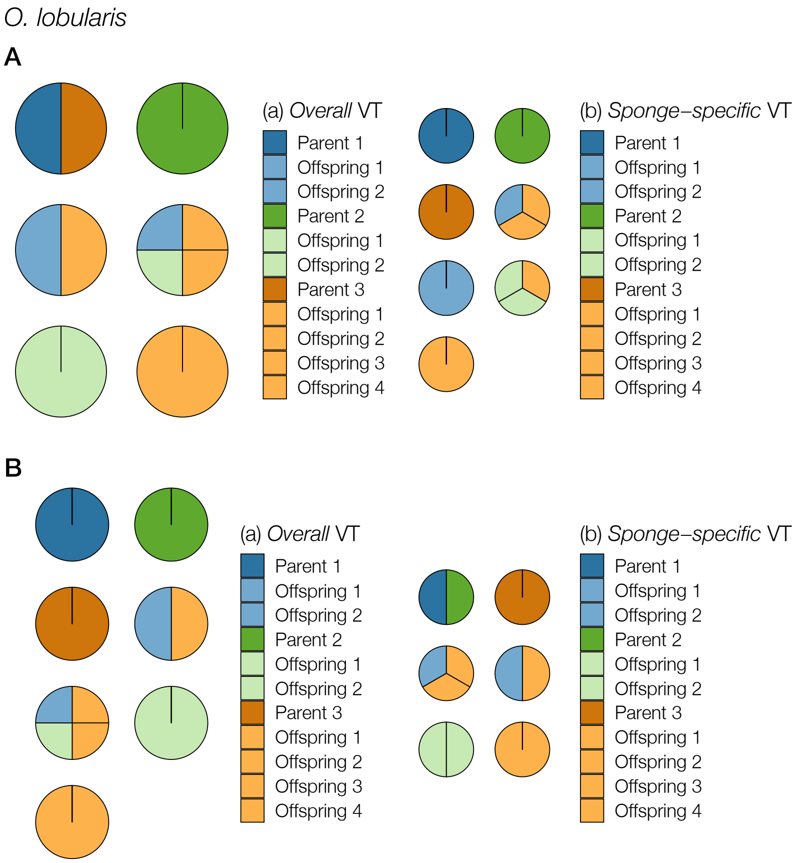


viii. See figure legend below

Figure S17: Host composition of the identified modules for (A) weighted and (B) unweighted networks constructed for each species individually. Panels (a) and (b) show modularity computed on networks corresponding to the *overall* and *sponge-specific* definition of vertical transmission, respectively. If parents and offspring harbor the same microbes, then parents and offspring are expected to form perfect compartments (i.e., modules only containing parents and their offspring). While weighted modularity weights edges by the relative abundance of ASV across hosts, unweighted modularity only considers the presence of ASVs in hosts. Each color represent one parent and its offspring, and circles represent the identified modules (the number of circles represent the number of identified modules). In the case of perfect parent-offspring compartments, there would be three unicolored modules. Panels i-viii corresponds to host species: *A. aerophoba, I. oros; I. fasciculata; C. crambe; C. viridis; D. avara; H. columella*; and *O. lobularis*.

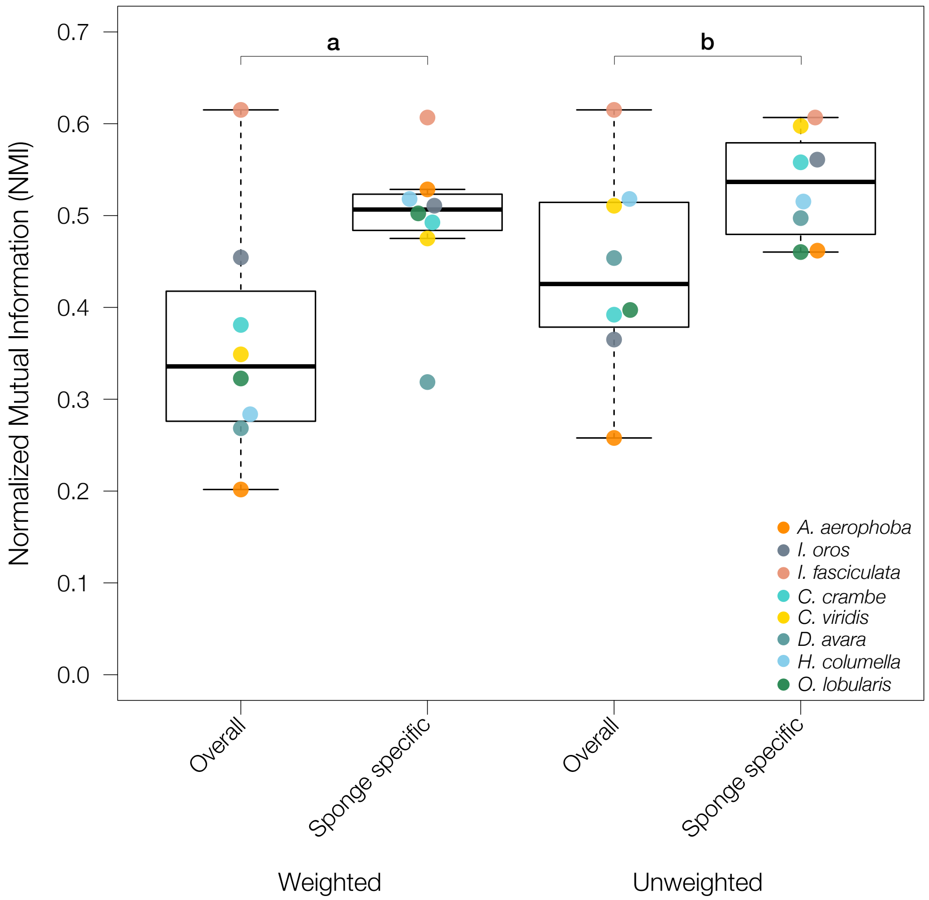


Figure S18: Boxplots for Normalized Mutual Information (NMI) criterion calculated for (a) weighted and (b) unweighted networks only containing conspecific hosts and their ASVs for *overall* and *sponge-specific* vertical transmission. Each colored dot corresponds to one sponge species. NMI ranges between 0 and 1, where 0 indicates complete dissimilarity between expected and observed modules, thus values closer to 1 corresponds to host species whose networks contain modules, and these modules contain nodes corresponding to parents and offspring.

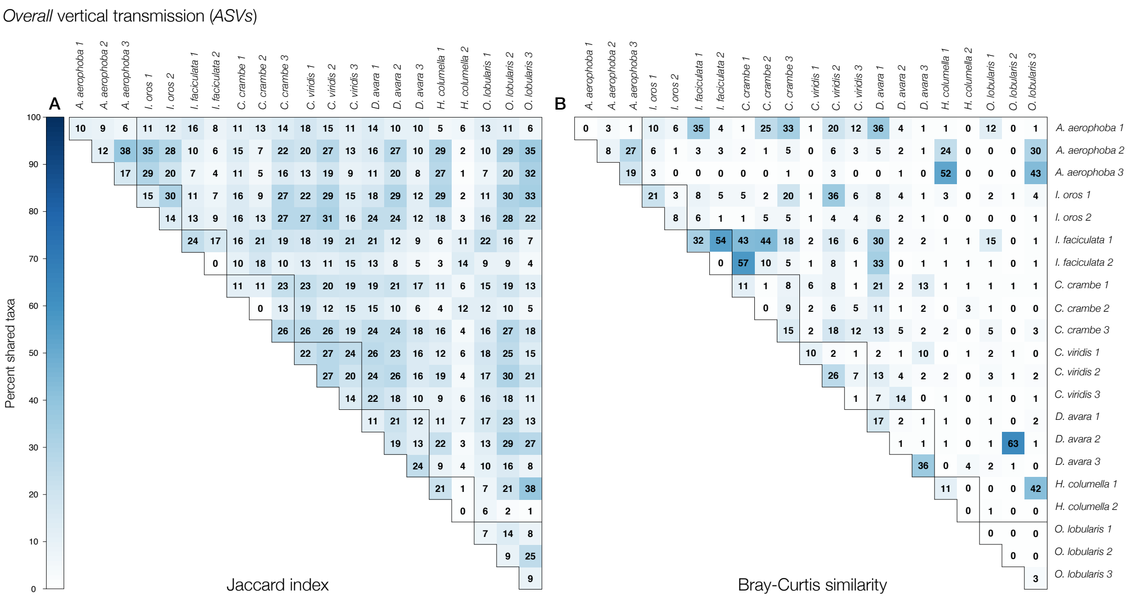


Figure S19: Similarity of *overall* vertically transmitted ASVs among all the offspring from to any given pair of adults. The figure shows similarity both within and between host species calculated as the (A) Jaccard index, or (B) Bray-Curtis similarity. In both panels, the diagonal corresponds to the average similarity among siblings (i.e., offspring from the same adult). While the Jaccard index calculates similarity between two samples based on the presence-absence of taxa, Bray-Curtis similarity also weights the coefficient by the number of individuals of each taxon. By converting the similarity coefficients to percents, their values range between 0% (no taxa shared; white), and 100% (all taxa shared; dark blue).

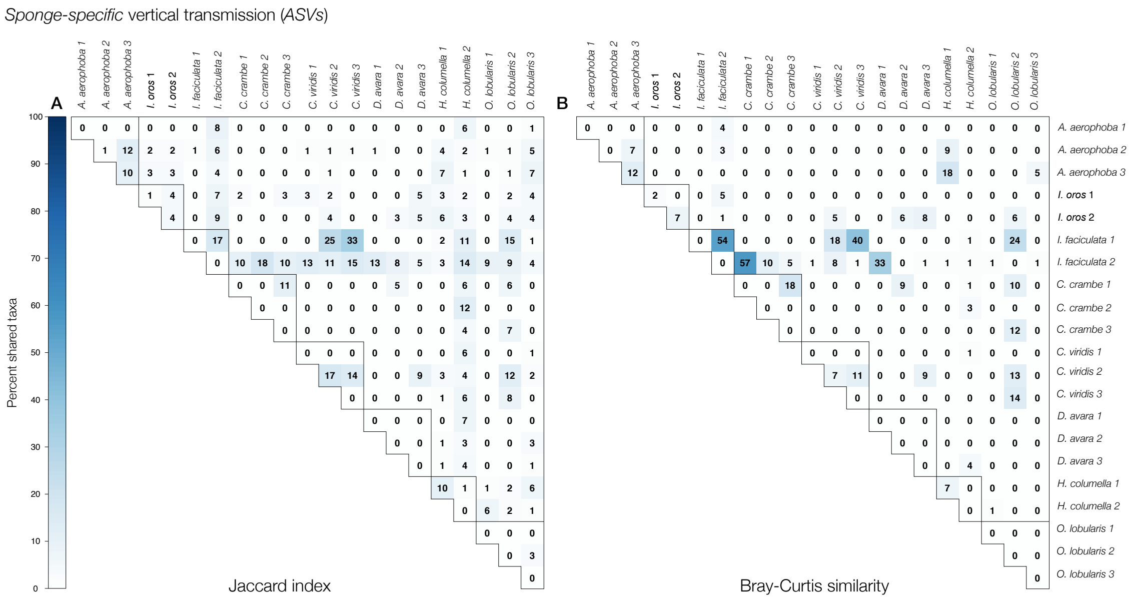


Figure S20: Similarity of *sponge-specific* vertically transmitted ASVs among all the offspring from to any given pair of adults. The figure shows similarity both within and between host species calculated as the (A) Jaccard index, or (B) Bray-Curtis similarity. In both panels, the diagonal corresponds to the average similarity among siblings (i.e., offspring from the same adult). While the Jaccard index calculates similarity between two samples based on the presence-absence of taxa, Bray-Curtis similarity also weights the coefficient by the number of individuals of each taxon. By converting the similarity coefficients to percents, their values range between 0% (no taxa shared; white), and 100% (all taxa shared; dark blue).

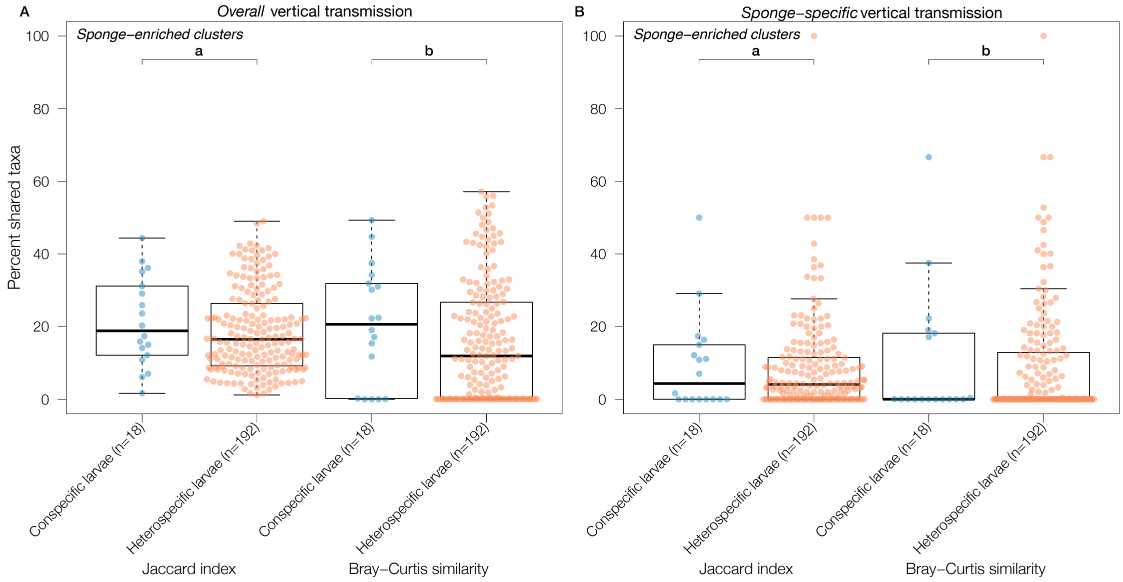


Figure S21: Percent shared (A) *overall* and (B) *sponge-specific* vertically transmitted *sponge-enriched clusters* among offspring from all possible combinations of adults calculated as either the (a) Jaccard index (see Figure S22 A and Figure S23 A), or (b) Bray-Curtis similarity (see Figure S22 B and Figure S23 B). Each dot represents all offspring from either (i) adults belonging to the same species (blue dots), or (ii) adults from different species (orange dots). While the Jaccard index calculates similarity between two samples based on the presence-absence of taxa, Bray-Curtis similarity also weights taxa by their relative abundance. For *overall* vertical transmission (A), conspecific larvae shared, on average, 21.3% (Jaccard) and 20.4% (Bray-Curtis) of the ASVs, whereas heterospecific larvae shared, on average, 18.6% (Jaccard) and 15.6% (Bray-Curtis) of the ASVs (P>0.1). For *sponge-specific* vertical transmission (B), parents and offspring shared, on average, 9.5% (Jaccard) and 10.1% (Bray-Curtis) of the ASVs, whereas non-parental conspecific adults and larvae shared, on average, 8.6% (Jaccard) and 8.5% (Bray-Curtis) of the ASVs (P>0.1). Note that the group *conspecific larvae* has much lesser number of observations (n) compared to the group *heterospecific larvae*.

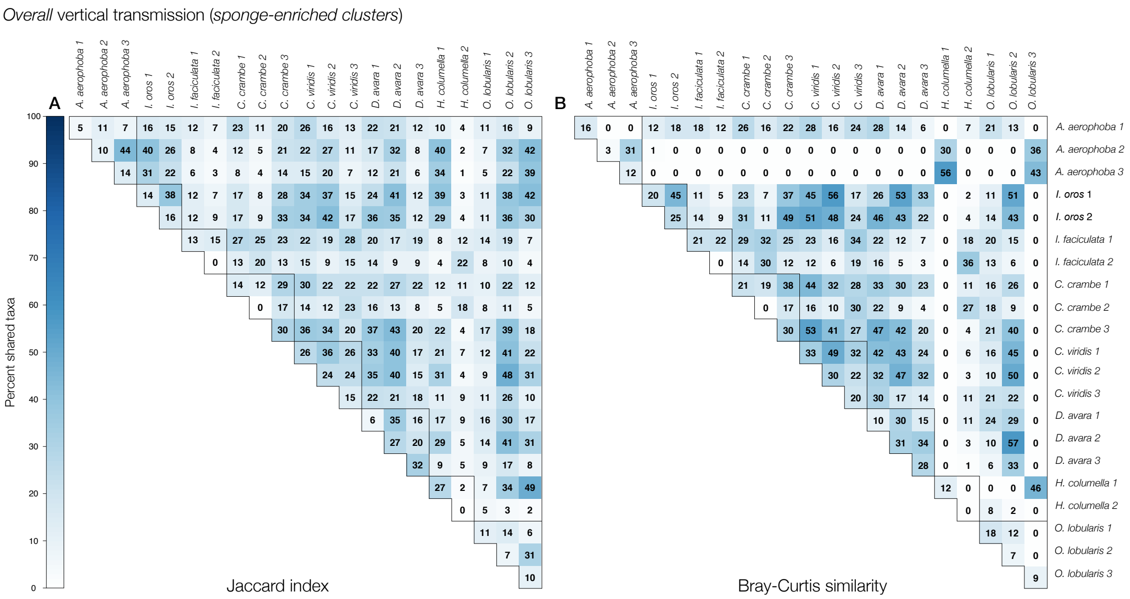


Figure S22: Similarity of *overall* vertically transmitted *sponge-enriched clusters* among all the offspring from to any given pair of adults. The figure shows similarity both within and between host species calculated as the (A) Jaccard index, or (B) Bray-Curtis similarity. In both panels, the diagonal corresponds to the average similarity among siblings (i.e., offspring from the same adult). While the Jaccard index calculates similarity between two samples based on the presence-absence of taxa, Bray-Curtis similarity also weights the coefficient by the number of individuals of each taxon. By converting the similarity coefficients to percents, their values range between 0% (no taxa shared; white), and 100% (all taxa shared; dark blue).

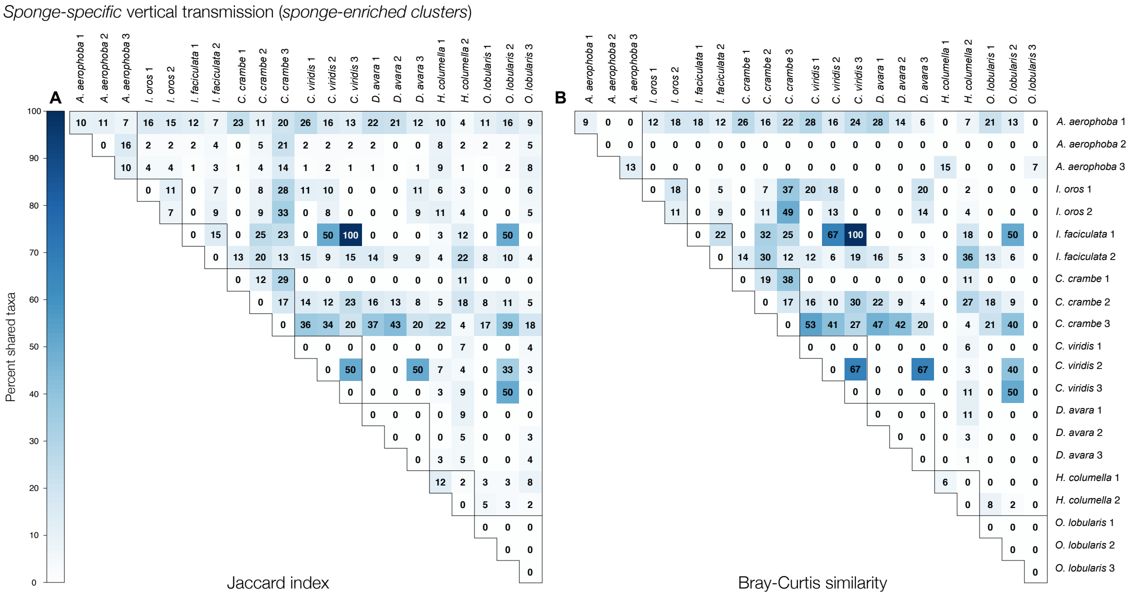


Figure S23: Similarity of *sponge-specific* vertically transmitted *sponge-enriched clusters* among all the offspring from to any given pair of adults. The figure shows similarity both within and between host species calculated as the (A) Jaccard index, or (B) Bray-Curtis similarity. In both panels, the diagonal corresponds to the average similarity among siblings (i.e., offspring from the same adult). While the Jaccard index calculates similarity between two samples based on the presence-absence of taxa, Bray-Curtis similarity also weights the coefficient by the number of individuals of each taxon. By converting the similarity coefficients to percents, their values range between 0% (no taxa shared; white), and 100% (all taxa shared; dark blue).

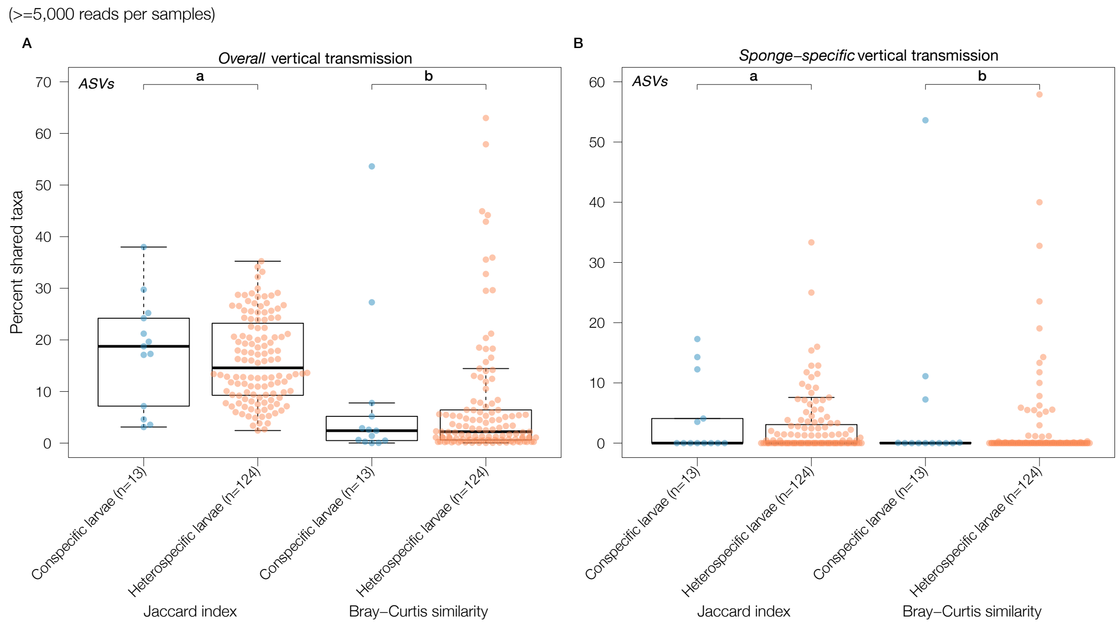


Figure S24: The percent of shared (A) *overall* and (B) *sponge-specific* vertically transmitted ASVs for samples with at least 5,000 sequence reads among offspring from all possible combinations of adults, calculated as either the (a) Jaccard index, or (b) Bray-Curtis similarity. Each dot represents all offspring from either (i) adults belonging to the same species (blue dots), or (ii) adults from different species (orange dots). While the Jaccard index calculates similarity between two samples based on the presence-absence of taxa, Bray-Curtis similarity weights taxa by their relative abundance. For *overall* vertical transmission (A), conspecific larvae shared, on average, 17.7% (Jaccard) and 8.0% (Bray-Curtis) of the ASVs, whereas heterospecific larvae shared, on average, 16.1% (Jaccard) and 7.1% (Bray-Curtis) of the ASVs (Jaccard: $\Delta$=1.11, 95% CI [-4.43,8.27], Mann-Whitney U=856, P>0.1; Bray-Curtis: $\Delta$=-0.22, 95% CI [-2.32,1.50], Mann-Whitney U=741, P>0.1). For *sponge-specific* vertical transmission (B), parents and offspring shared, on average, 4.0% (Jaccard) and 5.5% (Bray-Curtis) of the ASVs, whereas non-parental conspecific adults and larvae shared, on average, 2.6% (Jaccard) and 2.2% (Bray-Curtis) of the ASVs (Jaccard: $\Delta$=0.00, 95% CI [-0.00,0.20], Mann-Whitney U=824, P>0.1; Bray-Curtis: $\Delta$=-0.00, 95% CI [-0.00,0.00], Mann-Whitney U=801, P>0.1).

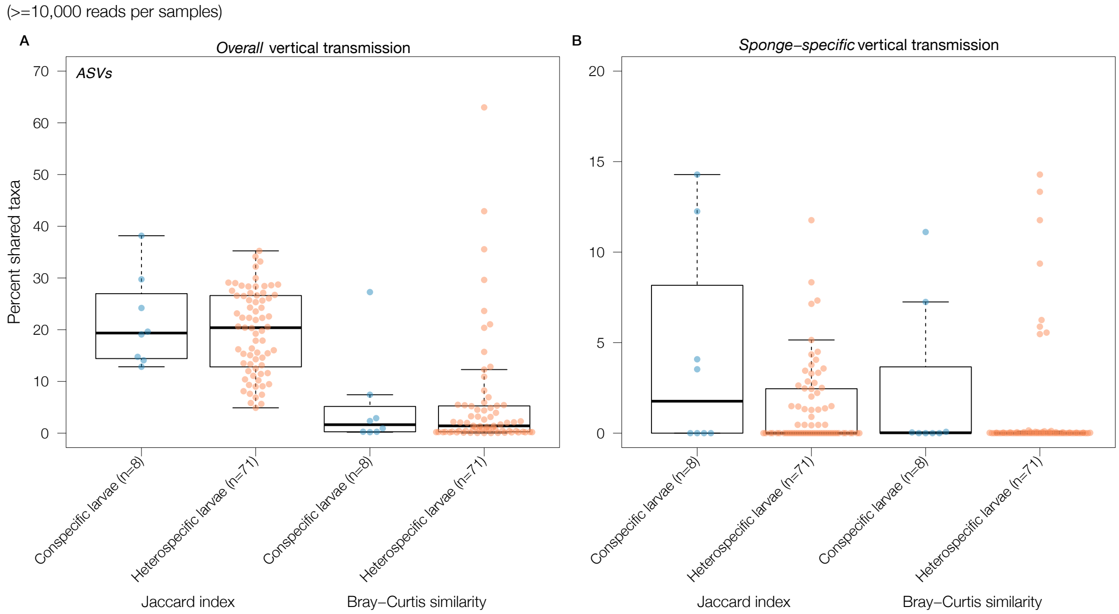


Figure S25: The percent of shared (A) *overall* and (B) *sponge-specific* vertically transmitted ASVs for samples with at least 10,000 sequence reads among offspring from all possible combinations of adults, calculated as either the (a) Jaccard index, or (b) Bray-Curtis similarity. Each dot represents all offspring from either (i) adults belonging to the same species (blue dots), or (ii) adults from different species (orange dots). While the Jaccard index calculates similarity between two samples based on the presence-absence of taxa, Bray-Curtis similarity weights taxa by their relative abundance. For *overall* vertical transmission (A), conspecific larvae shared, on average, 21.6% (Jaccard) and 5.2% (Bray-Curtis) of the ASVs, whereas heterospecific larvae shared, on average, 19.6% (Jaccard) and 5.6% (Bray-Curtis) of the ASVs (Jaccard: $\Delta$=1.60, 95% CI [-5.77,8.44], Mann-Whitney U=309, P>0.1; Bray-Curtis: $\Delta$=0.05, 95% CI [-2.10,2.15], Mann-Whitney U=295, P>0.1). For *sponge-specific* vertical transmission (B), parents and offspring shared, on average, 4.3% (Jaccard) and 2.3% (Bray-Curtis) of the ASVs, whereas non-parental conspecific adults and larvae shared, on average, 1.4% (Jaccard) and 1.0% (Bray-Curtis) of the ASVs (Jaccard: $\Delta$=0.00, 95% CI [-0.00,4.08], Mann-Whitney U=341, P>0.1; Bray-Curtis: $\Delta$=0.00, 95% CI [-0.01,0.07], Mann-Whitney U=319, P>0.1).

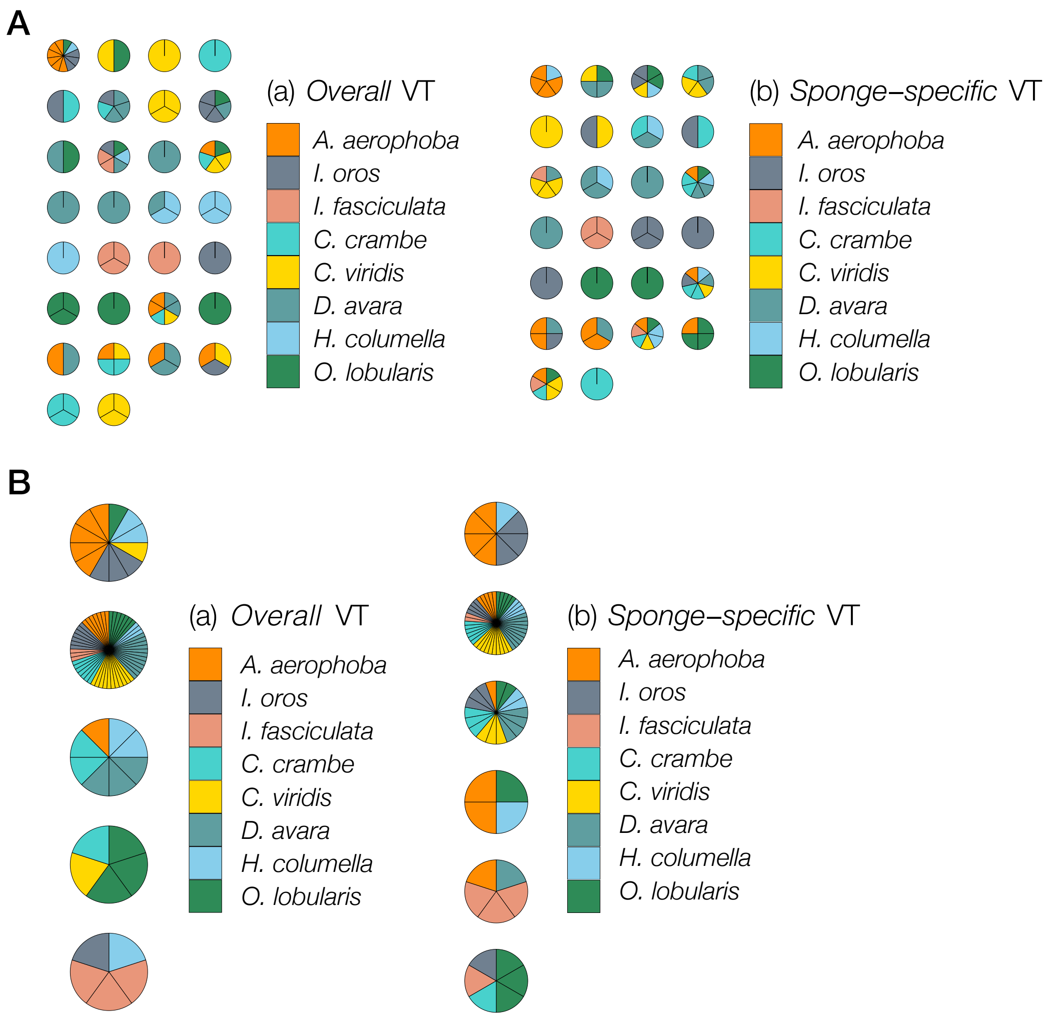


Figure S26: Host composition of the identified modules for a (A) weighted and (A) unweighted network containing all hosts (i.e., adults and larvae). If conspecific adults and larvae harbor the same microbes, and do not share those with other species, then the networks are expected to be organized in compartments consisting of conspecific adults and larvae only. While weighted modularity weights edges by the relative abundance of ASV across hosts, unweighted modularity only considers the presence of ASVs in hosts. Sponge species are represented by different colors. In the case of species-specific compartments, there would be eight unicolored modules.

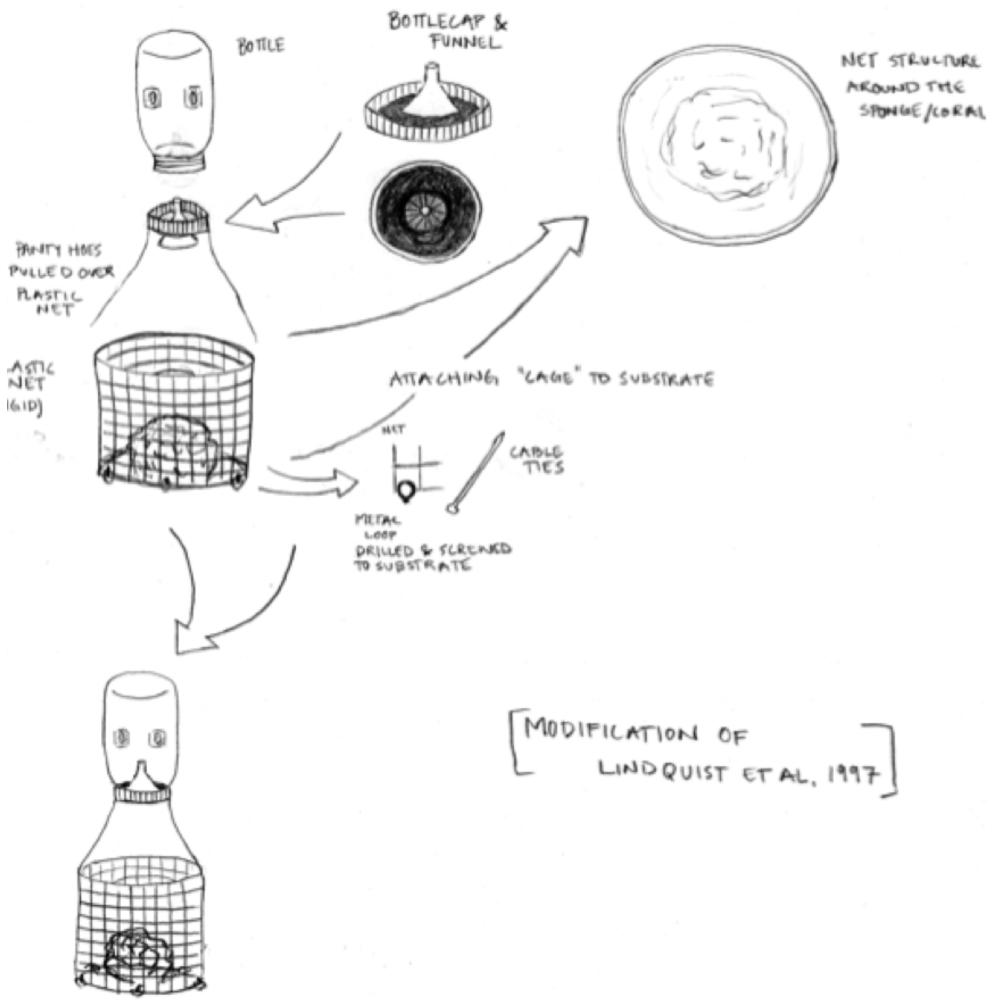


Figure S27: Sketch of the constructed traps that were used to capture dispersing larvae from adult sponges. Sketch by J.R.B.

### 1 Code Appendix

#### DADA2 Pipeline

Illumina-sequenced, single-read fastq files were processed and cleaned in R [87] using the default settings in DADA2 [88] to produce an amplicon sequence variant (ASV) table, and Silva (v128) [89] was used to create the ASV taxonomy.

```
library(dada2)
library(ShortRead)
path <- getwd() # change to the directory containing the fastq files after unzipping
print(path)
fns <- list.files(path, pattern="fastq.gz") # change if different file extensions
print(fns)
fastqs <- file.path(path, fns)
print(fastqs)

# Get sample names, assuming files named as so: SAMPLENAME_XXX.fastq
sample.names <- sapply(strsplit(basename(fns), "\\."), `[`, 1)
filt_path <- file.path(path, "filtered") # filtered files go into the filtered/ subdirectory
fns <- list.files(filt_path, pattern="fastq.gz")
sample.names <- sapply(strsplit(basename(fns), "\\."), `[`, 1)
print(sample.names)
if(!file_test("-d", filt_path)) dir.create(filt_path)
filts <- file.path(filt_path, paste0(sample.names,".fastq.gz"))

# Learn error rates
set.seed(100)
err <- learnErrors(filts, nreads = 2e6, multithread=TRUE, randomize=TRUE)

# Infer sequence variants
derep <- derepFastq(filts, verbose=TRUE)
names(derep) <- sample.names
dds <- dada(derep, err=err, multithread=TRUE, pool=TRUE)

# Construct sequence table and write to disk
seqtab <- makeSequenceTable(dds)
```

```r
seqtab.nochim <- removeBimeraDenovo(seqtab, method="consensus", multithread=TRUE) # remove chimeras
saveRDS(seqtab.nochim, paste0(filt_path,"/seqtab.nochim.RDS")) # change to the path where you want sequence table saved
getN <- function(x) sum(getUniques(x))
track <- cbind(out, sapply(dds, getN), rowSums(seqtab), rowSums(seqtab.nochim))
colnames(track) <- c("input", "filtered", "denoised", "tabled", "nonchim")
rownames(track) <- sample.names
saveRDS(track, paste0(filt_path,"/track_seqs.RDS")) # change to where you want sequence table saved

# Assign taxonomy
print("assign taxonomy")
seqtab.nochim <- readRDS("filtered/seqtab.nochim.RDS")
taxa <- assignTaxonomy(seqtab.nochim, "silva_nr_v128_train_set.fa",
        multithread=TRUE, verbose=TRUE, minBoot=80)
saveRDS(taxa, "taxa.RDS")
```

#### Initial filtering with Phyloseq

The `Phyloseq R` package [90] was used to filter out sequences classifying to *Archaea* and *Eukaryota*. We also removed singleton ASVs, and phyla that occurred in less than two samples. While the original analyzed dataset contained samples with at least 1,000 sequences, we also investigate how sequencing depth affected the results; we therefore created two additional datasets only containing samples with $\geq$ 5,000, and $\geq$ 10,000 sequences.

```r
library(phyloseq)
# Read in data
seqtab.nochim <- readRDS("seqtab.nochim.RDS") #ASV table
track <- readRDS("track_seqs.RDS") #sample data
taxa <- readRDS("taxa.RDS") #taxonomy

ps <- phyloseq(otu_table(seqtab.nochim, taxa_are_rows=FALSE),
               sample_data(track),
               tax_table(taxa))

# Filtering
ps0 <- subset_taxa(ps, !is.na(Kingdom) & !Kingdom %in% c("","Archaea","Eukaryota"))
```

```r
ps0 <- subset_taxa(ps0, !Phylum %in% c("", "uncharacterized"))
table(tax_table(ps0)[, "Phylum"], exclude = NULL)

# Compute prevalence of each feature, store as data.frame
prevdf = apply(X = otu_table(ps0),
               MARGIN = ifelse(taxa_are_rows(ps0), yes = 1, no = 2),
               FUN = function(x){sum(x > 0)})
# Add taxonomy and total read counts to this data.frame
prevdf = data.frame(Prevalence = prevdf,
                    TotalAbundance = taxa_sums(ps0),
                    tax_table(ps0))

prevdf_tbl <- plyr::ddply(prevdf, "Phylum", function(df1){cbind(mean(df1$Prevalence),sum(
    df1$Prevalence))})

# Remove phyla occuring in <=2 samples
ps1 = subset_taxa(ps0, !Phylum %in% as.character(prevdf_tbl[prevdf_tbl$`2`<=2,]$Phylum))

# Subset to the remaining phyla
prevdf1 = subset(prevdf, Phylum %in% get_taxa_unique(ps1, "Phylum"))
prevdf1_tbl <- plyr::ddply(prevdf1, "Phylum", function(df1){cbind(mean(df1$Prevalence),sum(
    df1$Prevalence))})

# Remove singleton AVSs, i.e. any given ASV must be present in at least 2 samples
prevalenceThreshold = 2

# Execute prevalence filter, using `prune_taxa()` function
keepTaxa = rownames(prevdf1)[(prevdf1$Prevalence >= prevalenceThreshold)]

ps2 = prune_taxa(keepTaxa, ps1)
prevdf2 = subset(prevdf1, Phylum %in% get_taxa_unique(ps2, "Phylum"))

ggplot(prevdf2, aes(TotalAbundance, Prevalence / nsamples(ps2),color=Phylum)) +
  geom_point(size = 2, alpha = 0.7) + scale_x_log10() + xlab("Total Abundance") + ylab("
      Prevalence [Frac. Samples]") + facet_wrap(~Phylum) + theme(legend.position="none")

sample_data(ps2)$nonchim <- rowSums(otu_table(ps2))

```

```r
# Remove samples that has < 1000 sequences (original analysis)
ps2_1000 <- subset_samples(ps2, nonchim>=1000)

# Additional analysis to investigate read depth

# Remove samples that has < 5000 sequences
ps2_5000 <- subset_samples(ps2, nonchim>=5000)

# Remove samples that has < 10,000 sequences
ps2_10000 <- subset_samples(ps2, nonchim>=10000)
```
